# *MAPT* regulates autophagic-lysosomal function and phagocytosis in human microglia

**DOI:** 10.64898/2026.08.27.747662

**Authors:** Kylie J. Schache, Rui Zhang, Audrey E. Street, Emma Starr, Jacob A. Marsh, David J. Kast, Sally Temple, Abhirami K. Iyer, Celeste M. Karch

## Abstract

Tauopathies are characterized by the accumulation and spread of pathogenic tau aggregates throughout the brain, a process that is increasingly recognized to involve not only neurons but also microglia. However, whether pathogenic *MAPT* directly alters microglial degradative capacity remains poorly understood. Here, using isogenic human induced pluripotent stem cell-derived microglia carrying the pathogenic *MAPT* IVS10+16 mutation, we identify tau as a regulator of microglial lysosomal function. *MAPT* IVS10+16 microglia exhibited coordinated suppression of lysosomal and autophagic pathways, reduced lysosomal protease abundance and activity, and impaired autophagosome-lysosome fusion. Mutant microglia also showed reduced uptake of extracellular tau aggregates, reduced tau accumulation in acidic compartments, and a blunted lysosomal response to proteopathic stress. Conversely, genetic loss of *MAPT* increased lysosomal degradative capacity and accumulation of extracellular tau aggregates within acidic compartments, supporting a cell-intrinsic role for endogenous tau in regulating microglial degradative function. Pharmacologic enhancement of the autophagy lysosome pathway in *MAPT* IVS10+16 microglia increased proteolytic activity and improved tau handling. Together, these findings reveal a reciprocal relationship between tau and microglial lysosome function and identify degradative capacity as a modifiable component of the microglial response to tau pathology.

## Introduction

Tauopathies are neurodegenerative diseases characterized by the progressive accumulation and spread of pathogenic tau aggregates throughout the brain^1^. Increasing evidence suggests that extracellular tau assemblies contribute to templated propagation of tau pathology and disease progression^2–7^, highlighting the importance of cellular pathways that regulate uptake, trafficking, and degradation of pathogenic tau species. Although neurons are the primary source of tau, multiple cell types contribute to the handling of extracellular tau aggregates, including microglia, the resident immune cells of the central nervous system^8^. Microglia dynamically respond to proteopathic stress, internalizing extracellular tau aggregates through phagocytosis and degradation via endolysosomal pathways^9–13^. However, failure of microglia-mediated degradation can lead to tau secretion and spread^9,10,14,15^, implicating microglial degradative function as a potential regulator of tau pathology.

Lysosomal and autophagic pathways are central to microglial function, supporting cargo degradation and proteostasis while regulating phagocytosis and immune signaling through the trafficking, turnover, and recycling of cell-surface receptors. Genetic studies implicate microglia-enriched genes that regulate endolysosomal function, including *TREM2*, *GRN*, and *TMEM106B*, in tauopathies and other neurodegenerative diseases^16–26^. Consistent with these genetic findings, studies of tauopathy patient tissues have identified alterations in autophagic and endolysosomal proteins, supporting disruption of degradative pathways in disease^27–33^. Studies in neuronal and immortalized cell models support a bidirectional relationship between tau pathology and degradative dysfunction, whereby pathogenic tau accumulates in endolysosomal compartments and impairs autophagic degradation, while impaired lysosomal function, in turn, promotes further tau accumulation and propagation^30,34–39^. Consistent with these findings, accumulation of tau and phosphorylated tau increases in mouse models of tauopathy upon disruption of autophagy-lysosome function *in vivo*^30,40–42^. Most studies linking tau to lysosomal dysfunction have focused on neuronal systems; therefore, the contribution of these pathways in microglia is not as well understood. Loss of microglial autophagy has been shown to exacerbate tau pathology in a mouse model^41^, demonstrating that microglial degradative pathways can influence tau pathology. However, whether pathogenic tau itself alters degradative function in microglia remains unknown.

Pathogenic *MAPT* mutations are sufficient to cause familial frontotemporal dementia with tau pathology (FTLD-tau), establishing that altered tau biology is a primary driver of disease^43^. The *MAPT* IVS10+16 mutation increases exon 10 inclusion and 4R tau expression, leading to FTLD-tau^43,44^. Prior work demonstrated that *MAPT* IVS10+16 induced pluripotent stem cell (iPSC)-derived microglia exhibit cell-autonomous changes in inflammatory states and reduced uptake of extracellular tau aggregates^45^, suggesting that pathogenic *MAPT* signaling alters microglial responses to tau. However, whether *MAPT* IVS10+16 disrupts autophagy-lysosome function and the capacity of microglia to mount an adaptive degradative response to proteopathic stress remains unknown.

Here, we used isogenic human iPSC-derived microglia to determine how *MAPT* IVS10+16 and endogenous tau influence autophagy-lysosome function and the processing of extracellular tau aggregates. We discovered that *MAPT* IVS10+16 impaired lysosomal proteolysis and autophagosome-lysosome fusion and reduced tau fibril processing, whereas genetic loss of *MAPT* and pharmacologic autophagy activation enhanced degradative function. These findings identify tau as a cell-intrinsic regulator of microglial degradative capacity and establish the autophagy-lysosome pathway as a modifiable component of microglial tau handling.

## Results

### MAPT IVS10+16 shifts microglial transcriptional states toward lysosomal dysfunction

To define the cell-autonomous effects of *MAPT* IVS10+16 in microglia, we utilized *MAPT* IVS10+16 and isogenic wild-type (WT) pairs derived from three independent donor iPSC backgrounds (**Figure 1A; Supplemental Table 1**). Two isogenic pairs were derived from patient lines heterozygous for *MAPT* IVS10+16 and CRISPR-corrected WT controls^45,46^. A third isogenic pair was generated by engineering one copy of the *MAPT* IVS10+16 mutation into a WT background^47^, providing a complementary genetic approach to distinguish mutation effects from donor-specific background variation (**Supplemental Figure 1**). iPSCs were differentiated through hematopoietic progenitor cells (HPCs) into microglia (iMG) using a growth factor-based protocol^45,48^ and analyzed at day 40 (**Figure 1A**). The resulting iMG were positive for CD45 and CD11b by flow cytometry (**Supplemental Figure 2A**) and TMEM119 and IBA1 by immunocytochemistry (**Figure 1B**; **Supplemental Figure 2C**). These findings confirmed successful microglial differentiation across genotypes, consistent with prior work showing that *MAPT* IVS10+16 does not prevent acquisition of microglial identity^45^.

**Figure 1.**
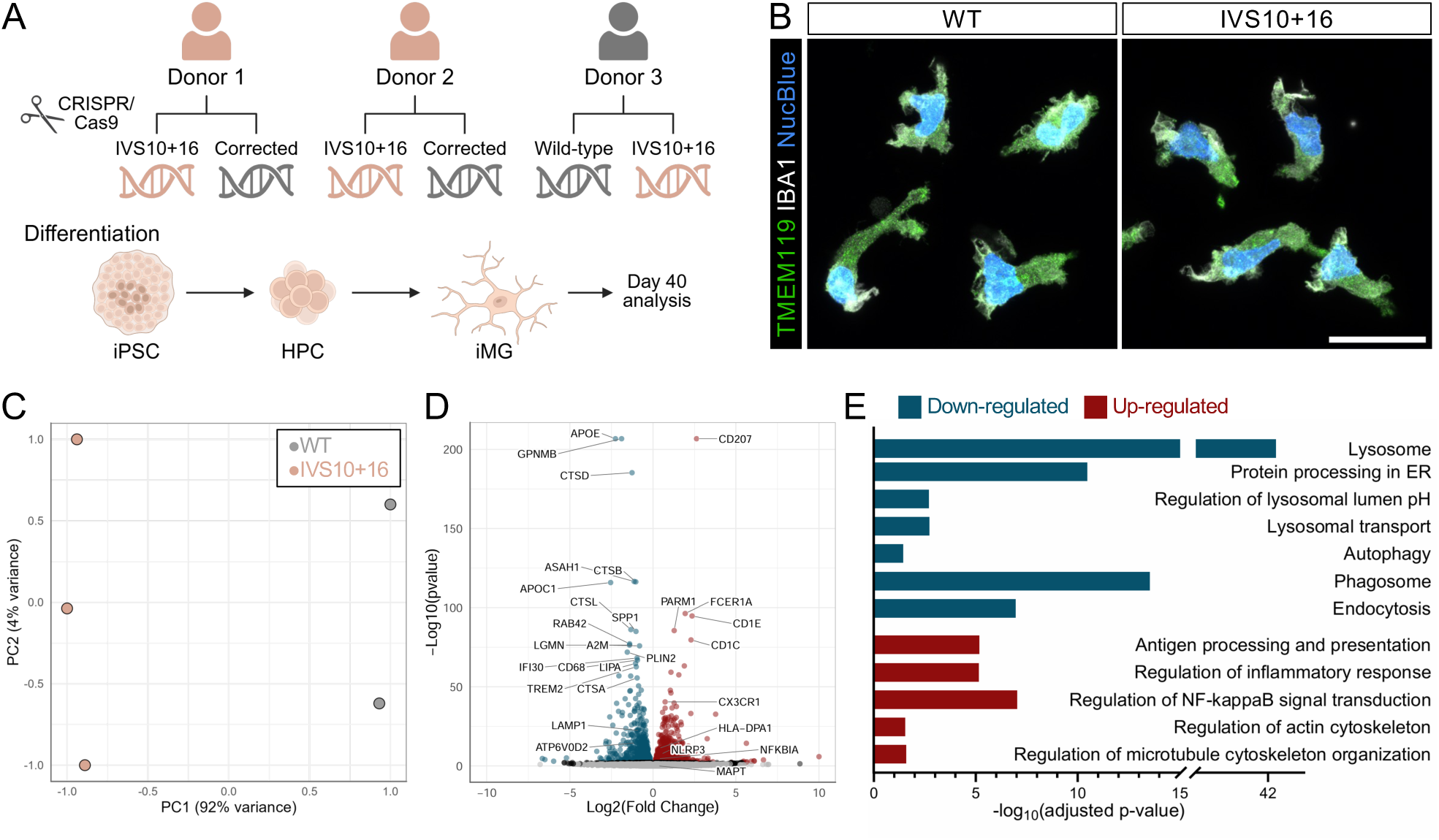
*MAPT* IVS10+16 disrupts expression of lysosome and phagocytosis gene networks in microglia. A. Schematic representing CRISPR editing strategy for generating three isogenic pairs of *MAPT* IVS10+16/WT and *MAPT* WT/WT iPSC lines and differentiation into microglia (iMG). B. Immunocytochemistry for microglia markers TMEM119 (green), IBA1 (white), and nuclei (blue). Representative max z projections from the GIH36 donor; additional donors in Supplemental Figure 2. Scale bar, 20μm. C-E. RNAseq and differential gene expression were performed on *MAPT* IVS10+16 iMG and isogenic controls from the GIH36 donor. WT, n=2; IVS10+16, n=3. C. Principal component analysis of the top 500 most variable genes using rlog-normalized counts in DESeq2. Grey, WT. Orange, IVS10+16. D. Volcano plot. Blue, significantly down-regulated genes (FDR p≤0.05; Log2FC<0). Red, significantly up-regulated genes (FDR p≤0.05; Log2FC>0). Black, p≤0.05. Grey, not significant. E. Enrichr pathway analysis of differentially expressed genes (FDR p≤0.05). Blue bars, down-regulated genes. Red bars, up-regulated genes.

To determine the extent to which the mutation alters microglial transcriptional states, we performed bulk RNA sequencing on *MAPT* IVS10+16 and isogenic control iMG. Principal component analysis (PCA) of the 500 most variable genes revealed distinct clustering between *MAPT* IVS10+16 and isogenic control iMG, suggesting that the *MAPT* mutation is sufficient to alter global transcriptional signatures in microglia (**Figure 1C**). Differential expression analysis identified broad transcriptional changes in *MAPT* IVS10+16 iMG relative to isogenic controls (**Figure 1D; Supplemental Table 2**). Pathway analyses on significantly differentially expressed genes (FDR≤0.05) revealed significant downregulation of pathways involved in lysosomal function, including lysosomal lumen pH, lysosomal transport, autophagy, phagosomes, and endocytosis in *MAPT* IVS10+16 iMG (**Figure 1E**). Consistent with these pathway-level changes, multiple lysosomal enzymes (e.g. *CTSD*, *CTSB*, *ASAH1*, *CTSL*, *IFI30*, *LIPA*, *CTSA*) were significantly downregulated in *MAPT* IVS10+16 iMG (FDR≤0.05; **Figure 1D-E; Supplemental Table 2**). We also observed significant upregulation of pathways supporting microglia immune and cytoskeletal functions, including antigen processing and presentation, NF-κB signaling, inflammatory responses, and actin and microtubule organization (**Figure 1E**). Together, these findings demonstrate a transcriptional shift in *MAPT* IVS10+16 microglia characterized by suppression of lysosomal and autophagy-related programs alongside increased immune and cytoskeletal pathways.

### Lysosome abundance and morphology are preserved in MAPT IVS10+16 microglia

Given the transcriptional repression of lysosomal and autophagy-related pathways in *MAPT* IVS10+16 iMG, we next sought to determine whether these changes were accompanied by alterations in lysosome abundance or morphology. Immunocytochemistry for the lysosomal marker LAMP1 revealed no overt differences in lysosomal morphometry between *MAPT* IVS10+16 and isogenic control iMG (**Figure 2A, Supplemental Figure 3A**). Quantification of LAMP1+ vesicle number, lysosomal volume, and overall LAMP1 intensity showed no significant differences between genotypes (**Figure 2A-C; Supplemental Figure 3A-D**). To further evaluate lysosomal protein levels, we performed immunoblotting for LAMP1 and LAMP2A, the most abundant transmembrane proteins on the lysosome^49,50^. LAMP1 and LAMP2A protein levels were comparable between *MAPT* IVS10+16 and isogenic control iMG (**Figure 2D-E; Supplemental Figure 3E-F**). Together, these findings suggest that lysosome abundance and morphology are largely preserved in *MAPT* IVS10+16 microglia despite transcriptional suppression of lysosomal and autophagic pathways.

**Figure 2.**
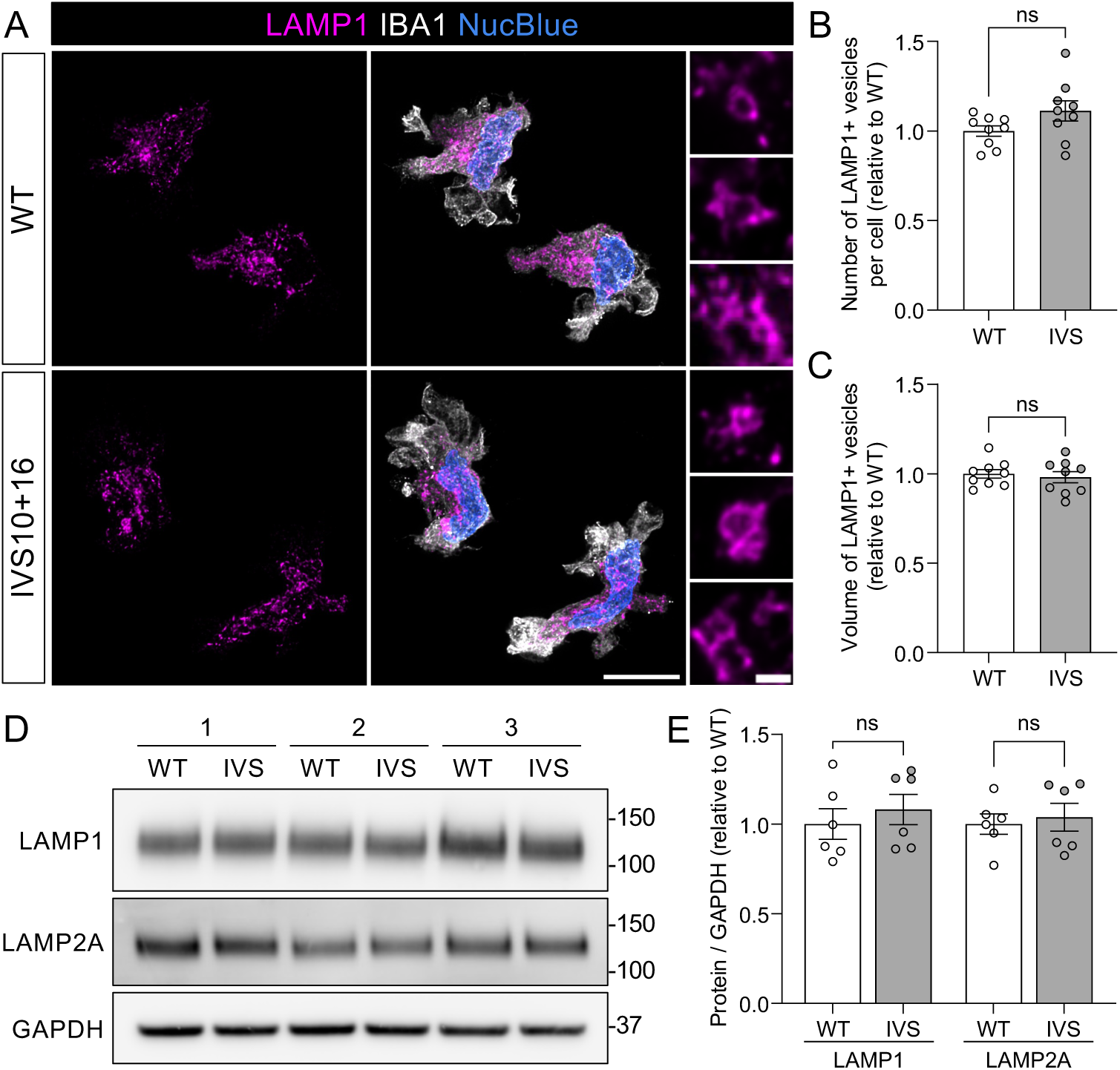
Lysosomal abundance and morphology are preserved in *MAPT* IVS10+16 microglia. A-C. Immunocytochemistry for LAMP1-positive vesicles. A. Representative max z projections of LAMP1 (magenta), IBA1 (white), and nuclei (blue). Large field scale bar, 10μm. Zoomed insets show the ring-shaped LAMP1-positive structures consistent with labeling limited to the vesicle membrane. Inset scale bar, 1μm. B. Quantification of LAMP1+ vesicle number per cell was performed in Imaris. Two-tailed unpaired t-test; p=0.0954. C. Quantification of LAMP1+ vesicle volume was performed in Imaris. Two-tailed unpaired t-test; p=0.6359. B-C. Data from three independent differentiations were normalized to WT within each differentiation. Each point represents data from three independent wells. WT, n=269 cells; IVS10+16, n=268. D-E. Whole cell lysates were analyzed for lysosomal membrane proteins by SDS-PAGE immunoblotting. D. Representative immunoblot for LAMP1, LAMP2A, and GAPDH (loading control) showing three independent differentiations. E. LAMP1 and LAMP2A protein levels quantified by densitometry, normalized to GAPDH, and expressed relative to WT controls. n=6 differentiations. Two-tailed paired t-test; p=0.1408 (LAMP1) and p=0.5033 (LAMP2A). Graphs represent mean ± SEM. ns, p>0.05. Data from GIH36 donor; additional donors in Supplemental Figure 3.

### MAPT IVS10+16 impairs lysosomal protease expression and degradative function

Although lysosome abundance and morphology were preserved in *MAPT* IVS10+16 iMG, transcriptomic analyses pointed to extensive suppression of lysosomal and autophagic pathways. We therefore examined whether *MAPT* IVS10+16 alters degradative machinery and function. We first analyzed expression of lysosomal enzymes spanning proteases, phosphatases, nucleases, glucosidases, sulfatases, and hydrolases involved in lipid and glycosaminoglycan metabolism^50,51^. Of the 62 lysosomal enzyme genes examined, 36 genes were significantly downregulated and 2 were significantly upregulated in *MAPT* IVS10+16 iMG relative to isogenic controls (FDR≤0.05; **Figure 3A; Supplemental Table 2**). Lysosomal enzyme genes were significantly enriched among differentially expressed genes compared with the expected frequency across the expressed transcriptome (Fisher’s exact test, p<0.0001; **Figure 3A**). These changes included multiple functional classes, such as proteases, phosphatases, nucleases, glucosidases, sulfatases, and enzymes involved in lipid and glycosaminoglycan catabolism (**Figure 3A**).

**Figure 3.**
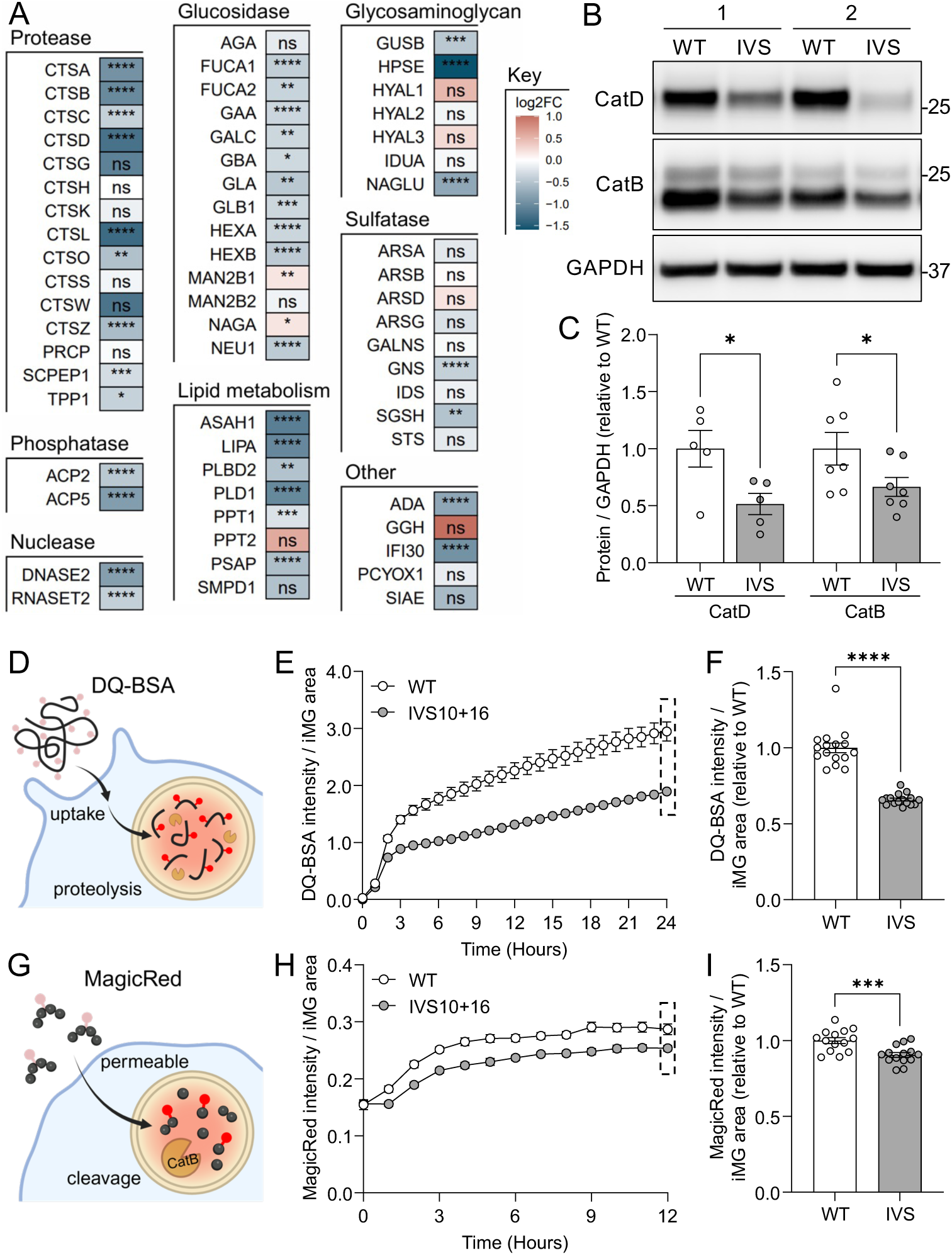
*MAPT* IVS10+16 disrupts lysosomal protease machinery and impairs degradative function in microglia. A. Heatmap of lysosomal enzyme genes differential expression in *MAPT* IVS10+16 iMG compared to isogenic controls. Genes are categorized based on the function of their substrate. Log2FC and FDR plotted for each gene. ns, FDR>0.05; *FDR≤0.05; **FDR≤0.01; ***FDR≤0.001; ****FDR≤0.0001. B-C. Whole cell lysates were analyzed for cathepsin proteins by SDS-PAGE immunoblotting. B. Representative immunoblot for cathepsin D, cathepsin B, and GAPDH (loading control) showing two independent differentiations. C. Cathepsin D and B protein levels quantified by densitometry, normalized to GAPDH, and expressed relative to WT controls. CatD, n=5 differentiations; CatB, n=7 differentiations. Two-tailed paired t-test (paired by differentiation); p=0.0160 (CatD) and p=0.0103 (CatB). D-F. To measure overall lysosomal proteolysis, iMG were treated with DQ-BSA (1μg/mL) and analyzed by Incucyte live cell imaging over 24hrs. D. Schematic. E. Quantification of DQ-BSA integrated intensity normalized to cell area over time from a representative differentiation. n=8 wells per genotype. F. Quantification of DQ-BSA at 24hrs. Data from two independent differentiations were normalized to WT within each differentiation. n=16 wells per genotype. Two-tailed unpaired t-test with Welch’s correction; p<0.0001. G-I. To evaluate cathepsin B proteolytic activity, iMG were treated with MagicRed (1:1000) and analyzed by Incucyte live cell imaging over 12hrs. G. Schematic. H. Quantification of MagicRed integrated intensity normalized to cell area over time from a representative differentiation. n=6 wells per genotype. I. Quantification of MagicRed at 12hrs. Data from two independent differentiations were normalized to WT within each differentiation. n=14 wells per genotype. Two-tailed unpaired t-test; p=0.0009. Graphs represent mean ± SEM. ns, p>0.05; *p≤0.05; ***p≤0.001; ****p≤0.0001. Data from GIH36 donor; additional donors in Supplemental Figure 4.

To determine whether these transcriptional changes extended to lysosomal protease abundance, we next examined cathepsin B and D protein levels in *MAPT* IVS10+16 iMG. Cathepsin proteases are synthesized as inactive precursors that undergo proteolytic maturation to generate active lysosomal enzymes^52,53^. Although the active form of cathepsin D is predicted to be approximately 34 kDa^52^, prior immunoblot studies commonly detect the mature enzyme between 25-28kDa^54–56^. Consistent with this, we detected cathepsin D in iMG at approximately 25 kDa (**Figure 3B, Supplemental Figure 4A**, uncropped blots in **Supplemental Figure 5A and 5C-D**). Cathepsin B is processed into two enzymatically active forms, the less active single chain and the more active heavy chain^53^. Immunoblotting in iMG detected both forms close to the expected size (**Figure 3B, Supplemental Figure 4C**, uncropped blots in **Supplemental Figure 5B, E-F**). *MAPT* IVS10+16 exhibited significantly reduced levels of mature cathepsin D and cathepsin B proteins relative to isogenic controls (**Figure 3B-C; Supplemental Figure 4A-D**).

We next asked whether reduced lysosomal enzyme expression was accompanied by impaired proteolytic function. Lysosomal proteolysis was assessed using DQ-BSA, a self-quenched substrate that fluoresces following lysosomal proteolysis (**Figure 3D**). *MAPT* IVS10+16 iMG exhibited reduced DQ-BSA fluorescence over time relative to isogenic controls (**Figure 3E**), with significantly reduced fluorescence observed at 24 hours (**Figure 3F; Supplemental Figure 4E**), indicating reduced lysosomal proteolytic capacity. We independently measured cathepsin B enzymatic activity using Magic Red, a membrane-permeable fluorogenic substrate containing a cathepsin B preferred cleavage sequence (**Figure 3G**). *MAPT* IVS10+16 iMG exhibited reduced Magic Red fluorescence over time relative to isogenic controls (**Figure 3H**), with significantly reduced fluorescence observed at 12 hours (**Figure 3I; Supplemental Figure 4F**). Together, these findings demonstrate that *MAPT* IVS10+16 produces coordinated reductions in lysosomal enzyme expression, mature cathepsin abundance, and lysosomal proteolytic function in human microglia.

### MAPT IVS10+16 impairs autophagic responses and autophagosome-lysosome convergence

Although *MAPT* IVS10+16 iMG exhibited impaired lysosomal protease expression and degradative function, it remained unknown whether autophagic pathways were also altered in mutant microglia. To determine whether lysosomal dysfunction extended to autophagic cargo delivery, we next examined autophagic responses and autophagosome-lysosome fusion in *MAPT* IVS10+16 iMG (**Figure 4A**). To assess autophagic vesicle dynamics, iMG were labeled with CYTO-ID, a live-cell fluorescent probe that labels autophagosomes and autolysosomes (**Figure 4A**). To induce autophagic responses, iMG were treated with rapamycin and chloroquine (Rap+CQ), which induces autophagic vesicle accumulation by stimulating autophagy while impairing lysosomal acidification and autophagosome-lysosomal fusion. Autophagic vesicle content was assessed via flow cytometry, and two-way ANOVA of CYTO-ID mean fluorescence intensity (MFI) identified significant effects of genotype and treatment (**Figure 4B**). Post-hoc analyses revealed significantly reduced CYTO-ID MFI in *MAPT* IVS10+16 iMG relative to isogenic controls under both vehicle-and Rap+CQ-treated conditions, suggesting an overall reduction in autophagic vesicle content or altered vesicle maturation in mutant microglia (**Figure 4B; Supplemental Figure 6A**). Rap+CQ treatment significantly increased CYTO-ID MFI in both genotypes relative to their respective vehicle-treated conditions, indicating that autophagic pathways remained responsive to pharmacologic stimulation despite the genotype-dependent deficit (**Figure 4B**). These findings pointed to an overall difference in autophagic vesicle content and prompted us to examine vesicle morphometry in greater detail.

**Figure 4.**
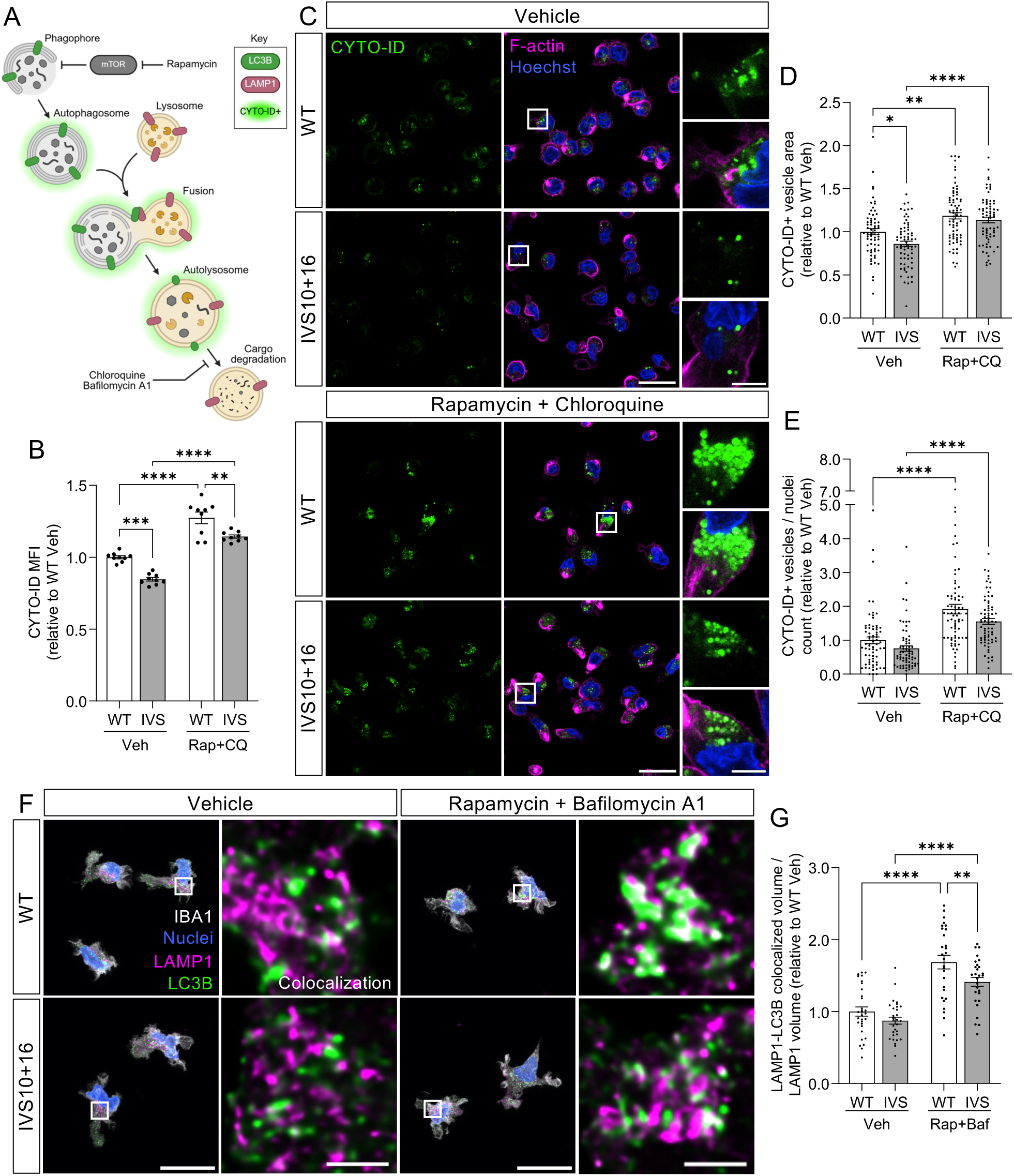
*MAPT* IVS10+16 impairs autophagy by disrupting autophagosome-lysosome fusion in microglia. A. Schematic. CYTO-ID (green glow) is a live cell dye which labels autophagic vesicles (autophagosomes and autolysosomes). LC3B (green) is a marker of autophagic vesicles. LAMP1 (magenta) is a marker of lysosomes. B-E. iMG were treated with rapamycin (500nM, Rap) and chloroquine (10μM, CQ) or DMSO control (Veh) for 6hrs and labeled with CYTO-ID. B. Flow cytometry. Quantification of CYTO-ID geometric mean fluorescent intensity (MFI) within the live cell population. Data from three independent differentiations were normalized to WT Veh within each differentiation. n=9 wells per group. Two-way ANOVA: genotype F(1, 32)=40.66, p<0.0001; treatment F(1, 32)=164.9, p<0.0001; interaction F(1, 32)=0.2767, p=0.6025. Tukey’s multiple comparisons test: WT Veh vs IVS Veh, p=0.0002; WT Veh vs WT Rap+CQ, p<0.0001; IVS Veh vs IVS Rap+CQ, p<0.0001; WT Rap+CQ vs IVS Rap+CQ, p=0.0013. C-E. Live cell confocal microscopy. C. Representative images of autophagic vesicles (CYTO-ID, green) with F-actin (SpyActin, magenta) and nuclei (Hoechst, blue). Large field scale bar, 25μm; zoom inset scale bar, 5μm. D. Quantification of CYTO-ID+ vesicle area expressed relative to WT Veh. Two-way ANOVA: genotype F(1, 273)=7.515, p=0.0065; treatment F(1, 273)=44.78, p<0.0001; interaction F(1, 273)=1.844, p=0.1756. Tukey’s multiple comparisons test: WT Veh vs IVS Veh, p=0.0240; WT Veh vs WT Rap+CQ, p=0.0012; IVS Veh vs IVS Rap+CQ, p<0.0001; WT Rap+CQ vs IVS Rap+CQ, p=0.7527. E. Quantification of CYTO-ID+ vesicle number normalized to nuclei per field and expressed relative to WT Veh. Two-way ANOVA: genotype F(1, 277)=8.170, p=0.0046; treatment F(1, 277)=66.36, p<0.0001; interaction F(1, 277)=0.3210, p=0.5715. Tukey’s multiple comparisons test: WT Veh vs IVS Veh, p=0.3767; WT Veh vs WT Rap+CQ, p<0.0001; IVS Veh vs IVS Rap+CQ, p<0.0001; WT Rap+CQ vs IVS Rap+CQ, p=0.0715. D-E. 20-26 fields were acquired per condition for three independent differentiations. Each point represents one field. Data were normalized to WT Veh within each differentiation. WT Veh, n=546 cells; IVS10+16 Veh, n=467; WT Rap+CQ, n=434; IVS10+16 Rap+CQ, n=484. F-G. To assess autophagosome-lysosome fusion, iMG were treated with rapamycin (1µM, Rap) and bafilomycin A1 (400nM, Baf) or DMSO (Veh) and analyzed by immunocytochemistry. F. Whole field, representative max z projections of LAMP1 (magenta), LC3B (green), IBA1 (white), and nuclei (blue). Scale bar, 20μm. Zoomed inset, representative single z slice images of LAMP1 and LC3B with co-localized area in white. Scale bar, 2μm. G. Quantification of LAMP1-LC3B co-localized volume normalized to total LAMP1 volume per field. Each point represents one field. Ten fields were acquired per group for three independent differentiations, with data normalized to WT Veh within each differentiation. WT Veh, n=78 cells; IVS10+16 Veh, n=76; WT Rap+Baf, n=73; IVS10+16 Rap+Baf, n=73. Two-way ANOVA: genotype F(1, 112)=8.512, p=0.0043; treatment F(1, 112)=79.48, p<0.0001; interaction F(1, 112)=1.135, p=0.2891. Fisher’s LSD test: WT Veh vs IVS Veh, p=0.1853; WT Veh vs WT Rap+Baf, p<0.0001; IVS Veh vs IVS Rap+Baf, p<0.0001; WT Rap+Baf vs IVS Rap+Baf, p=0.0066. Graphs represent mean ± SEM. ns, p>0.05; *p≤0.05; **p≤0.01; ***p≤0.001; ****p≤0.0001. Data from GIH36 donor; additional donors in Supplemental Figure 6.

To further assess autophagic vesicle properties, we used live cell confocal imaging to visualize CYTO-ID+ vesicles in iMG and quantified vesicle area (a proxy for vesicle size) and vesicle number (normalized to nuclei number per field; **Figure 4C-E**). Two-way ANOVA identified significant genotype and treatment effects on CYTO-ID+ vesicle area. Post-hoc analyses revealed that *MAPT* IVS10+16 iMG exhibited significantly reduced CYTO-ID+ vesicle area relative to isogenic controls in the vehicle-treated condition (**Figure 4D**). As expected, Rap+CQ treatment significantly increased vesicle area in both genotypes (**Figure 4D**). For CYTO-ID+ vesicle number, two-way ANOVA identified significant genotype and treatment effects (**Figure 4E**). Post-hoc multiple comparisons confirmed that Rap+CQ treatment significantly increased CYTO-ID+ vesicle number in both genotypes but did not identify significant genotype-dependent differences within individual treatment conditions (**Figure 4E**). Together, these findings reveal that *MAPT* IVS10+16 iMG exhibit reduced autophagic vesicle size compared to isogenic controls in vehicle-treated conditions without a consistent reduction in vesicle number.

Given the altered autophagic vesicle size observed in *MAPT* IVS10+16 iMG, we next examined autophagosome-lysosome fusion. Because microglia are challenging to efficiently transfect or transduce with conventional autophagic flux reporters, we assessed autophagosome-lysosome fusion by immunocytochemistry for LC3B, a marker of autophagosomes, and LAMP1, a marker of lysosomes (**Figure 4F**). LC3B and LAMP1 co-localization reflects the physical convergence of autophagosomes with lysosomes, a critical step required for cargo degradation and completion of autophagic flux. Two-way ANOVA identified that changes in LAMP1 and LC3B co-localization were associated with genotype and treatment (**Figure 4G**; **Supplemental Figure 6B**). Post hoc comparisons revealed that LAMP1-LC3B co-localization was similar between *MAPT* IVS10+16 and isogenic control iMG under basal conditions, suggesting that steady-state autophagosome-lysosome interactions are largely preserved in mutant microglia (**Figure 4G**; **Supplemental Figure 6B**). Following rapamycin and Bafilomycin A1 treatment, both genotypes exhibited increased LAMP1-LC3B co-localization compared to their respective vehicle controls. Interestingly, treated *MAPT* IVS10+16 iMG displayed significantly reduced co-localization relative to treated controls (**Figure 4G**; **Supplemental Figure 6B**). This reduction in co-localization under autophagic stress conditions suggests that *MAPT* IVS10+16 iMG are less capable of efficiently recruiting or fusing lysosomes with autophagosomes when autophagic demand is elevated. Together, these results suggest that *MAPT* IVS10+16 alters adaptive autophagic responses in human microglia and limits autophagosome-lysosome convergence during stress.

### MAPT IVS10+16 impairs tau uptake and adaptive lysosomal responses in microglia

A primary role of microglia in tauopathies is to phagocytose and degrade tau aggregates. Given our observation that *MAPT* IVS10+16 impaired lysosomal degradative function at baseline and stress-induced autophagosome-lysosome convergence in iMG, we next asked whether *MAPT* IVS10+16 alters the ability of microglia to process disease-relevant tau aggregates. Previous work demonstrated reduced uptake of myelin and tau preformed fibrils (PFFs) in *MAPT* IVS10+16 iMG^45^. To define the transcriptional response to tau aggregates, we performed RNA sequencing on *MAPT* IVS10+16 and isogenic control iMG following tau PFF treatment. PCA revealed a clear transcriptional shift following tau PFF treatment in both genotypes (**Supplemental Figure 7A**). Differential expression analysis similarly identified broad transcriptional alterations in iMG following tau PFF treatment in both *MAPT* IVS10+16 and control iMG (**Supplemental Figure 7B-E; Supplemental Tables 4-5**). Pathway analysis on significantly differentially expressed genes (FDR≤0.05) identified enrichment of immune activation, phagocytosis, and lysosomal function in both genotypes following tau PFF treatment compared to vehicle-treated controls (**Supplemental Figure 7B-E**), consistent with a shared transcriptional response to extracellular tau that engages pathways involved in cargo uptake and lysosomal processing. However, direct comparison of tau PFF-treated *MAPT* IVS10+16 with control iMG revealed broad transcriptional differences between genotypes (**Figure 5A-B; Supplemental Figure 7A; Supplemental Table 3**). Tau PFF-treated *MAPT* IVS10+16 exhibited reduced enrichment of lysosomal, proteolytic, phagosomal, endocytic, lipid transport, antigen processing and presentation, and ER stress-response pathways, together with increased inflammatory, NF-κB, and cytoskeletal programs compared to control microglia (**Figure 5C**). Thus, although both genotypes mount a transcriptional response to tau PFF exposure, *MAPT* IVS10+16 microglia exhibit a relative suppression of degradative and trafficking pathways and a shift toward inflammatory and cytoskeletal programs compared to *MAPT* WT microglia.

**Figure 5.**
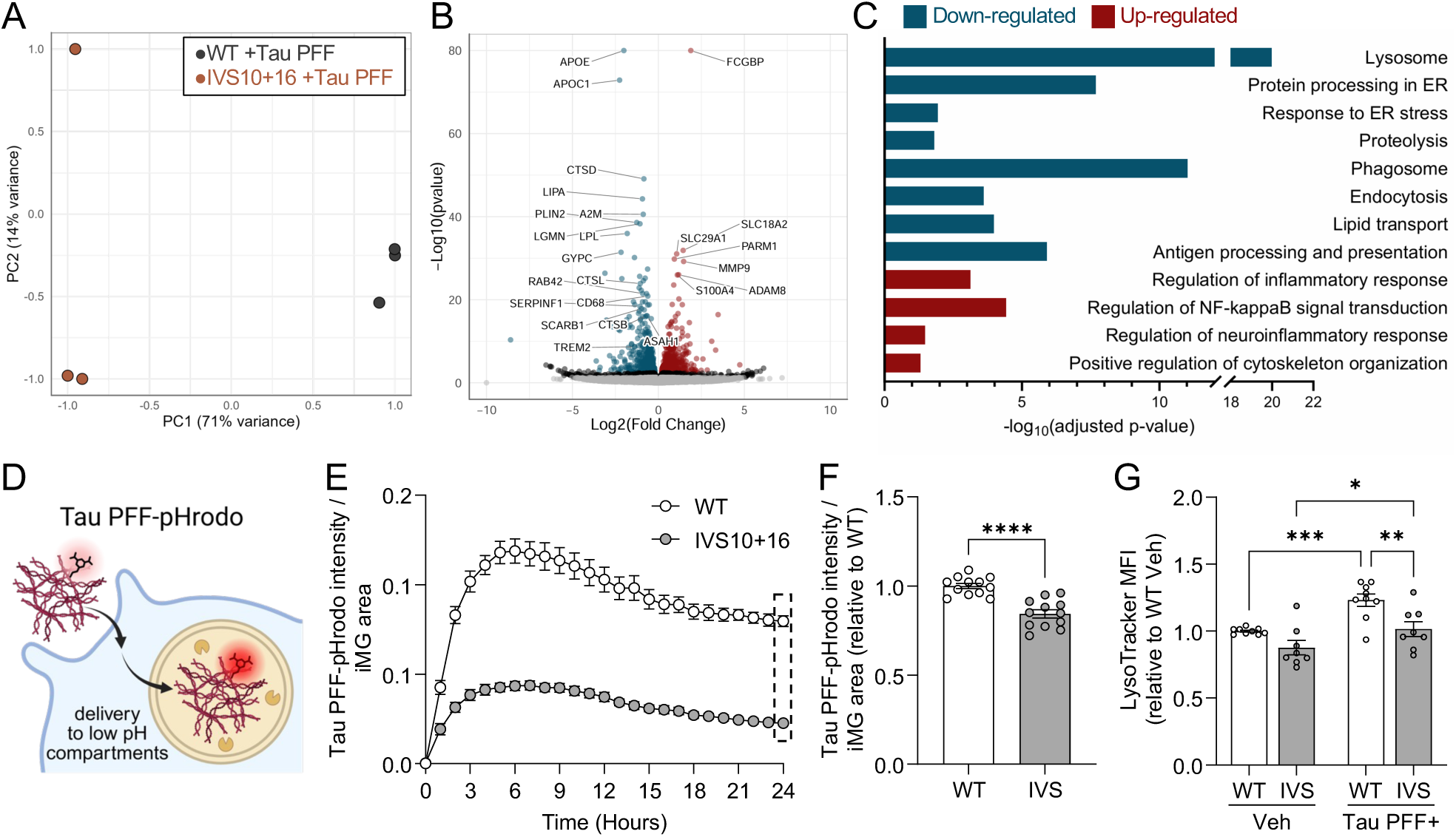
*MAPT* IVS10+16 impairs microglial processing of tau aggregates and adaptive lysosomal responses. A-C. *MAPT* IVS10+16 and isogenic control iMG were treated with tau preformed fibrils (tau PFF, 50nM for 24hrs) and analyzed by RNAseq. Samples represent different wells from a single differentiation. n=3 wells per genotype. A. Principal component analysis of the top 500 most variable genes using rlog-normalized counts in DESeq2. Grey, WT. Orange, IVS10+16. B. Volcano plot. Y-axis truncated at 80; APOE and FCGBP plotted at limit for visualization. Blue, significantly down-regulated genes (FDR p≤0.05; Log2FC<0). Red, significantly up-regulated genes (FDR p≤0.05; Log2FC>0). Black, p≤0.05. Grey, not significant. C. Enrichr pathway analysis of differentially expressed genes (FDR p≤0.05). Red bars, up-regulated genes. Blue bars, down-regulated genes. D-F. iMG were treated with tau PFF-pHrodo (250nM), which fluoresces in acidic environments, and analyzed by Incucyte live cell imaging over 24hrs. D. Schematic. E. Quantification of tau PFF-pHrodo integrated intensity normalized to cell area over time from a representative differentiation of the KOLF donor. n=6 wells per genotype. F. Quantification of tau PFF-pHrodo at 24hrs. Data from two independent differentiations were normalized to WT within each differentiation. n=12 wells per genotype. Two-tailed unpaired t-test; p<0.0001. G. iMG were treated with tau PFF-ATTO488 (500nM) for 24hrs, labeled with LysoTracker for 30mins, and analyzed by flow cytometry. Quantification of LysoTracker geometric mean fluorescent intensity (MFI) within the live cell population for Veh-treated samples and within the Tau PFF+ population for tau-treated samples. Data from three independent differentiations were normalized to WT within each differentiation. WT Veh, n=9 wells; IVS10+16 Veh, n=8; WT Tau PFF, n=9; IVS10+16 Tau PFF, n=8. Two-way ANOVA: genotype F(1, 30)=15.13, p=0.0005; treatment F(1, 30)=17.93, p=0.0002; interaction F(1, 30)=1.123, p=0.2978. Fisher’s LSD test: WT Veh vs IVS Veh, p=0.0545; WT Veh vs WT Tau, p=0.0006; IVS Veh vs IVS Tau, p=0.0371; WT Tau vs IVS Tau, p=0.0015. Graphs represent mean ± SEM. *p≤0.05; **p≤0.01; ***p≤0.001; ****p≤0.0001. Unless otherwise indicated, data from GIH36 donor; additional donors in Supplemental Figure 8.

To determine whether these transcriptional differences following tau PFF exposure were accompanied by altered tau processing, we first assessed uptake of fluorescently labeled tau fibrils. *MAPT* IVS10+16 and isogenic WT iMG were treated with ATTO488-labeled tau PFFs for 24 hours, and tau uptake was measured using flow cytometry. Consistent with prior findings^45^, all three *MAPT* IVS10+16 lines exhibited significantly reduced uptake of fluorescently labeled tau aggregates relative to isogenic controls (**Supplemental Figure 7F-G**). We next asked whether *MAPT* IVS10+16 also impairs delivery of internalized tau to acidic compartments. To address this, we used pHrodo-conjugated tau PFFs, which exhibit increased fluorescence upon delivery to acidic organelles such as lysosomes (**Figure 5D**). Live-cell imaging demonstrated significantly reduced pHrodo-tau fluorescence over time in *MAPT* IVS10+16 iMG relative to controls (**Figure 5E**), with significantly reduced fluorescence observed at 24 hours (**Figure 5F; Supplemental Figure 8A**). Because pHrodo fluorescence depends on tau uptake, delivery to acidic compartments, and acidification of the compartment, reduced pHrodo-tau PFF signal cannot distinguish among defects in these processes. Together with the independent reduction in ATTO488-tau uptake, these findings are consistent with impaired tau uptake and trafficking to lysosomes, potentially compounded by reduced lysosomal acidification in *MAPT* IVS10+16 iMG.

Lysosomal biogenesis and acidification are critical adaptive responses that enable microglia to degrade pathogenic protein aggregates; thus, we next asked whether *MAPT* IVS10+16 alters the lysosomal response to tau proteopathic stress. Following tau PFF exposure, we quantified acidified compartments using LysoTracker staining (**Figure 5G; Supplemental Figure 8B**). Two-way ANOVA identified significant effects of genotype and treatment on LysoTracker mean fluorescence intensity (MFI) (**Figure 5G; Supplemental Figure 8B**). Post-hoc analyses revealed tau PFF treatment significantly increased LysoTracker MFI in both *MAPT* IVS10+16 and isogenic control iMG (**Figure 5G; Supplemental Figure 8B**), consistent with a lysosomal response to extracellular tau and the induction of lysosomal transcriptional responses after tau PFF uptake (**Supplemental Figure 7C,E**). By post-hoc comparisons, LysoTracker MFI was significantly reduced in *MAPT* IVS10+16 compared with the isogenic control iMG in vehicle-treated conditions in one donor background but was not reproduced across donors. In contrast, following tau PFF treatment, *MAPT* IVS10+16 iMG exhibited significantly reduced LysoTracker MFI relative to controls across multiple donor backgrounds (**Figure 5G; Supplemental Figure 8B**). Thus, although mutant microglia retain the capacity to increase acidic lysosomal compartments in response to tau PFFs, mutant microglia ultimately exhibit a reduced level of lysosomal compartments compared to control microglia following tau PFF exposure. Together with the reduced tau uptake and pHrodo-tau signal, these findings demonstrate that *MAPT* IVS10+16 impairs microglial processing of extracellular tau aggregates and limits the adaptive lysosomal responses to proteopathic stress.

### Tau loss enhances lysosomal degradative function and tau aggregate processing in human microglia

Given the widespread impacts of the *MAPT* IVS10+16 mutation on microglial lysosomal function, we next asked whether endogenous tau plays a cell-intrinsic regulatory role in these processes. To determine whether endogenous tau contributes to regulation of lysosomal function in human microglia, we used a *MAPT* knockout (KO) iPSC line generated by CRISPR editing, followed by differentiation into iMG (**Figure 6A**; see methods for details). RT-PCR analyses confirmed loss of *MAPT* expression in the *MAPT* KO iMG compared to the isogenic *MAPT* WT control iMG (**Supplemental Figure 1D**). We first examined whether tau loss alters lysosomal degradative capacity using DQ-BSA live-cell imaging. Relative to isogenic control iMG, *MAPT* KO iMG exhibited a progressive increase in DQ-BSA fluorescence over time (**Figure 6B**), with significantly elevated fluorescence observed at 24 hours (**Figure 6C**). Thus, in contrast to the reduced proteolytic activity observed with *MAPT* IVS10+16, loss of endogenous *MAPT* enhanced lysosomal degradative capacity in human microglia.

**Figure 6.**
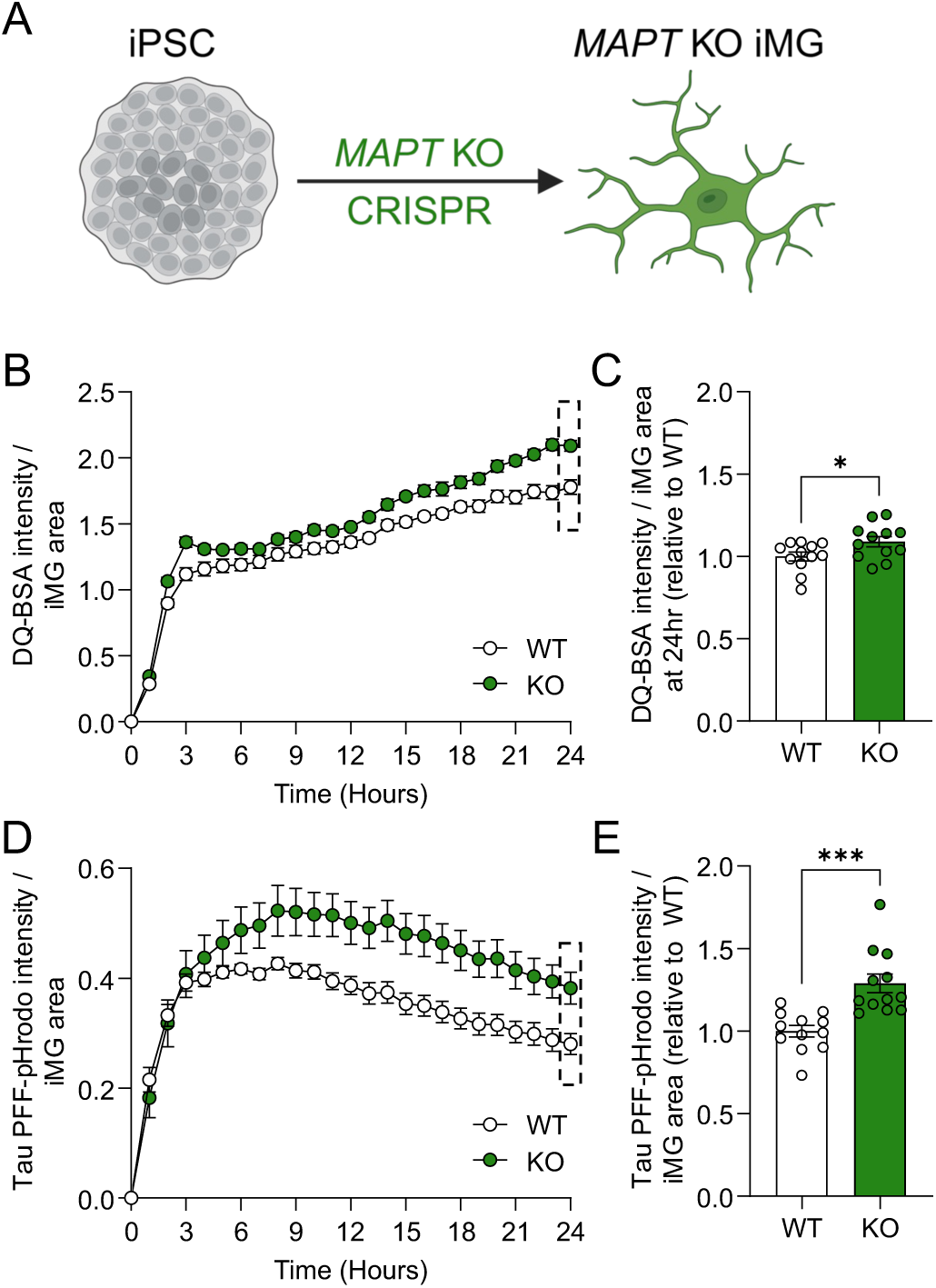
*MAPT* loss enhances lysosomal degradative function and tau aggregate processing in iMG. A. Schematic of *MAPT* KO engineering and differentiation. B-E. *MAPT* KO and isogenic control iMG were treated with fluorogenic substrates and analyzed by Incucyte live cell imaging over 24hrs. B. Quantification of DQ-BSA (1µg/uL) integrated intensity normalized to cell area over time from a representative differentiation. n=6 wells per genotype. C. Quantification of DQ-BSA at 24hrs. Data from two independent differentiations were normalized to WT within each differentiation. n=12 wells per genotype. Unpaired two-tailed t-test; p=0.0351. D. Quantification of tau PFF-pHrodo (250nM) integrated intensity normalized to cell area over time from a representative differentiation. n=6 wells per genotype. E. Quantification of tau PFF-pHrodo at 24hrs. Data from two independent differentiations were normalized to WT within each differentiation. n=12 wells per genotype. Unpaired two-tailed t-test; p=0.0003. Graphs represent mean ± SEM. *p≤0.05; ***p≤0.001.

We next asked whether this increase in degradative activity was accompanied by altered processing of extracellular tau aggregates by monitoring uptake and lysosomal delivery of pHrodo-conjugated tau PFFs. *MAPT* KO iMG exhibited increased pHrodo-tau fluorescence relative to isogenic controls (**Figure 6D**), with significantly elevated fluorescence observed at 24 hours (**Figure 6E**), suggesting that loss of *MAPT* enhances the uptake and delivery of extracellular tau PFFs to acidic degradative compartments. Together, these findings support a role for endogenous tau in constraining microglial lysosomal degradative function and tau aggregate processing.

### Rapamycin enhances lysosomal degradative function and tau processing in MAPT IVS10+16 microglia

Given the lysosomal degradative deficits and altered responses to proteopathic stress in *MAPT* IVS10+16 iMG, we asked whether pharmacological enhancement of the autophagy lysosome pathway could improve these phenotypes in mutant microglia. Rapamycin is a well-established inhibitor of mechanistic target of rapamycin complex 1 (mTORC1), a central negative regulator of autophagy and lysosomal biogenesis^57–59^. Inhibition of mTORC1 by rapamycin promotes autophagic flux and stimulates cellular degradative pathways through activation of transcriptional and metabolic programs that support lysosome-dependent clearance^57^. *MAPT* IVS10+16 iMG were treated with rapamycin (100nM) for 4 hours prior to assessment of lysosomal degradative activity and tau processing (**Figure 7A**). We first examined the effect of rapamycin on lysosomal proteolytic activity using DQ-BSA live-cell imaging. Consistent with our prior findings, vehicle-treated *MAPT* IVS10+16 iMG exhibited significantly reduced DQ-BSA fluorescence relative to vehicle-treated isogenic controls (**Figure 7B-D**). Rapamycin treatment increased DQ-BSA fluorescence in *MAPT* IVS10+16 iMG across the imaging period (**Figure 7B**), with significantly increased fluorescence observed at 12 hours relative to vehicle-treated *MAPT* IVS10+16 iMG (**Figure 7C-D**). The magnitude of the effect of rapamycin on protease activity varied across donor backgrounds (**Figure 7C-D**). These findings demonstrate that pharmacologic activation of the autophagy-lysosome pathway enhances lysosomal degradative function in *MAPT* IVS10+16 microglia.

**Figure 7.**
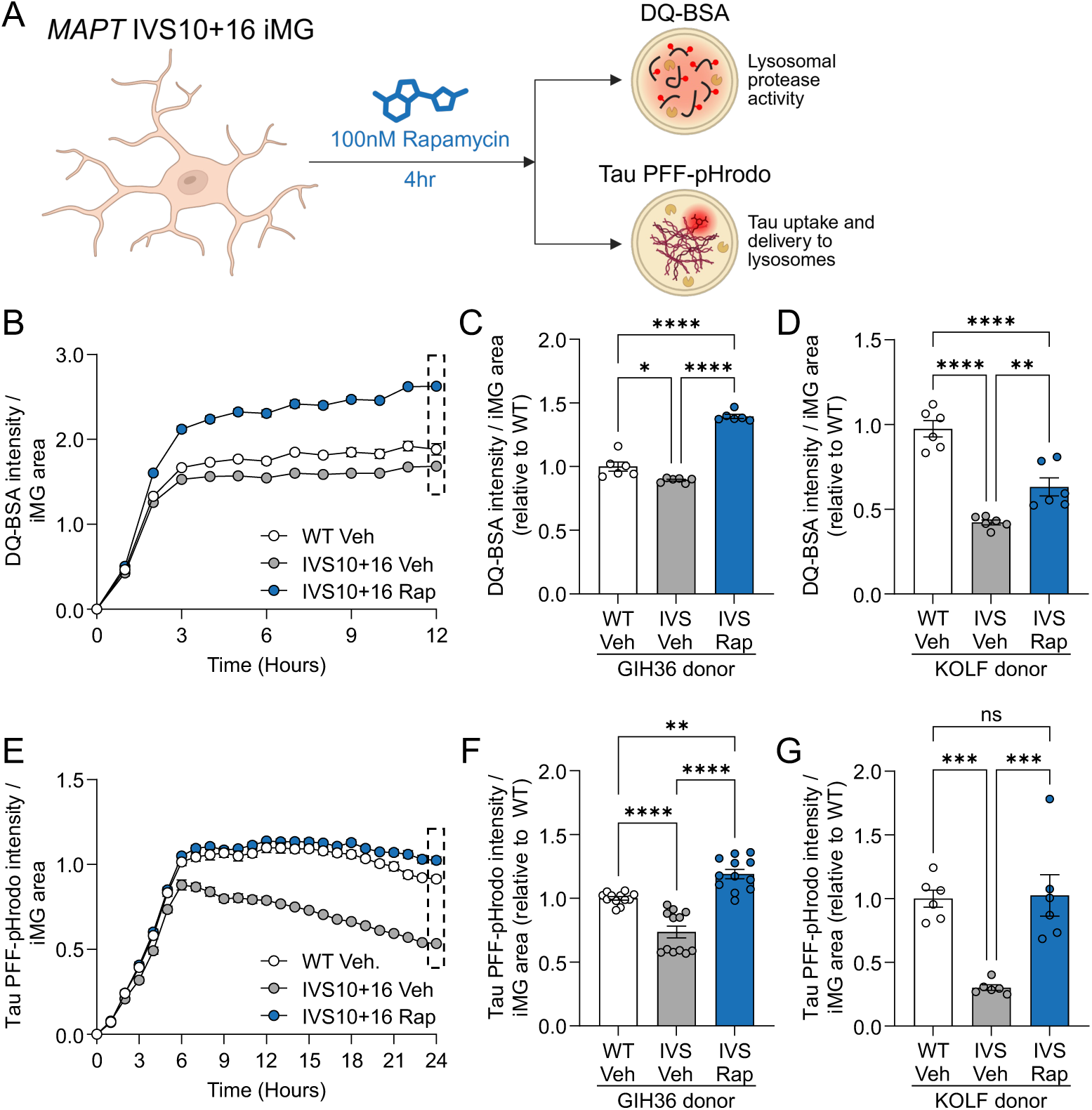
Rapamycin enhances lysosomal function and tau processing in *MAPT* IVS10+16 microglia. iMG were treated with rapamycin (100nM, Rap), an autophagy stimulator, or DMSO control (Veh) for 4hrs then treated with fluorogenic substrates and analyzed by Incucyte live cell imaging. A. Schematic. B-D. iMG were treated with DQ-BSA (1μg/mL) and imaged for 12hrs. B. Quantification of DQ-BSA integrated intensity normalized to cell area over time from a representative GIH36 differentiation. n=6 wells per group. C-D. Quantification of DQ-BSA at 12hrs. Data from one differentiation normalized to WT Veh. n=6 wells per group. C. GIH36 donor. Ordinary one-way ANOVA [F(2, 15)=131.3, p<0.0001] followed by Tukey’s multiple comparisons test: WT Veh vs IVS Veh, p=0.0137; IVS Veh vs IVS Rap, p<0.0001; WT Veh vs IVS Rap, p<0.0001. D. KOLF2.1J donor. Ordinary one-way ANOVA [F(2, 15)=44.30, p<0.0001] followed by Tukey’s multiple comparisons test: WT Veh vs IVS Veh, p<0.0001; IVS Veh vs IVS Rap, p=0.0080; WT Veh vs IVS Rap, p<0.0001. E-G. iMG were treated with tau PFF-pHrodo (250nM) and imaged for 24hrs. E. Tau PFF-pHrodo intensity normalized to cell area over time from one representative GIH36 differentiation. n=6 wells per group. F-G. Quantification of tau PFF-pHrodo at 24hrs. F. GIH36 donor. Data from two independent differentiations were normalized to WT Veh within each differentiation. n=12 wells per group. Ordinary one-way ANOVA [F(2, 33)=42.43, p<0.0001] followed by Tukey’s multiple comparisons test: WT Veh vs IVS Veh, p<0.0001; IVS Veh vs IVS Rap, p<0.0001; WT Veh vs IVS Rap, p=0.0014. G. KOLF donor. Data from one differentiation normalized to WT Veh. n=6 wells per group. Ordinary one-way ANOVA [F(2, 15)=16.04, p=0.0002] followed by Tukey’s multiple comparisons test. WT Veh vs IVS Veh, p=0.0006; IVS Veh vs IVS Rap, p=0.0004; WT Veh vs IVS Rap, p=0.9836. Graphs represent mean ± SEM. ns, p>0.05; *p≤0.05; **p≤0.01; ***p≤0.001; ****p≤0.0001.

Because microglia rely on coordinated endolysosomal and autophagic pathways to internalize, traffic, and degrade extracellular protein aggregates, we tested whether rapamycin-mediated enhancement of lysosomal function could improve processing of extracellular tau aggregates. To test this, we monitored pHrodo-conjugated tau PFFs fluorescence over time (**Figure 7E-G**). Vehicle-treated *MAPT* IVS10+16 iMG exhibited significantly reduced pHrodo-tau fluorescence relative to vehicle-treated isogenic controls, consistent with reduced delivery of tau PFFs to acidic degradative compartments (**Figure 7E-G**). Rapamycin treatment increased pHrodo-tau fluorescence in *MAPT* IVS10+16 iMG relative to vehicle-treated mutants (**Figure 7E**), with significantly increased fluorescence observed at 24 hours relative to vehicle-treated *MAPT* IVS10+16 iMG (**Figure 7F-G**). Following rapamycin treatment, pHrodo-tau signal in *MAPT* IVS10+16 iMG was restored to (**Figure 7G**) or exceeded WT levels (**Figure 7F**) depending on the donor background. Together with the increase in DQ-BSA proteolysis, these findings demonstrate that pharmacologic enhancement of the autophagy-lysosome pathway improves both lysosomal degradative capacity and delivery of extracellular tau aggregates to acidic compartments in *MAPT* IVS10+16 microglia.

## Discussion

Here, we describe a previously unrecognized role for tau in regulating the degradative state of human microglia, extending the effects of tau beyond neuronal proteostasis and suggesting that altered microglial lysosomal function may influence how extracellular tau is handled in tauopathy (**Figure 8**). *MAPT* IVS10+16 impaired microglial degradative function without overtly altering lysosomal abundance or morphology. Mutant microglia exhibited suppressed lysosomal protease expression, reduced degradative activity, limited autophagosome-lysosome fusion, and impaired microglial processing of extracellular tau aggregates. Loss of *MAPT* increased degradative activity and tau aggregate processing, revealing a role for tau in microglial lysosomal function. Pharmacological enhancement of lysosomal and autophagic pathways with rapamycin restored degradative function and tau processing in *MAPT* IVS10+16 microglia. Together, our findings establish lysosomal dysfunction as a central mechanism linking *MAPT* IVS10+16 to impaired microglial handling of pathogenic tau aggregates and suggest that modulation of lysosomal pathways may represent a therapeutic strategy for tauopathies.

**Figure 8.**
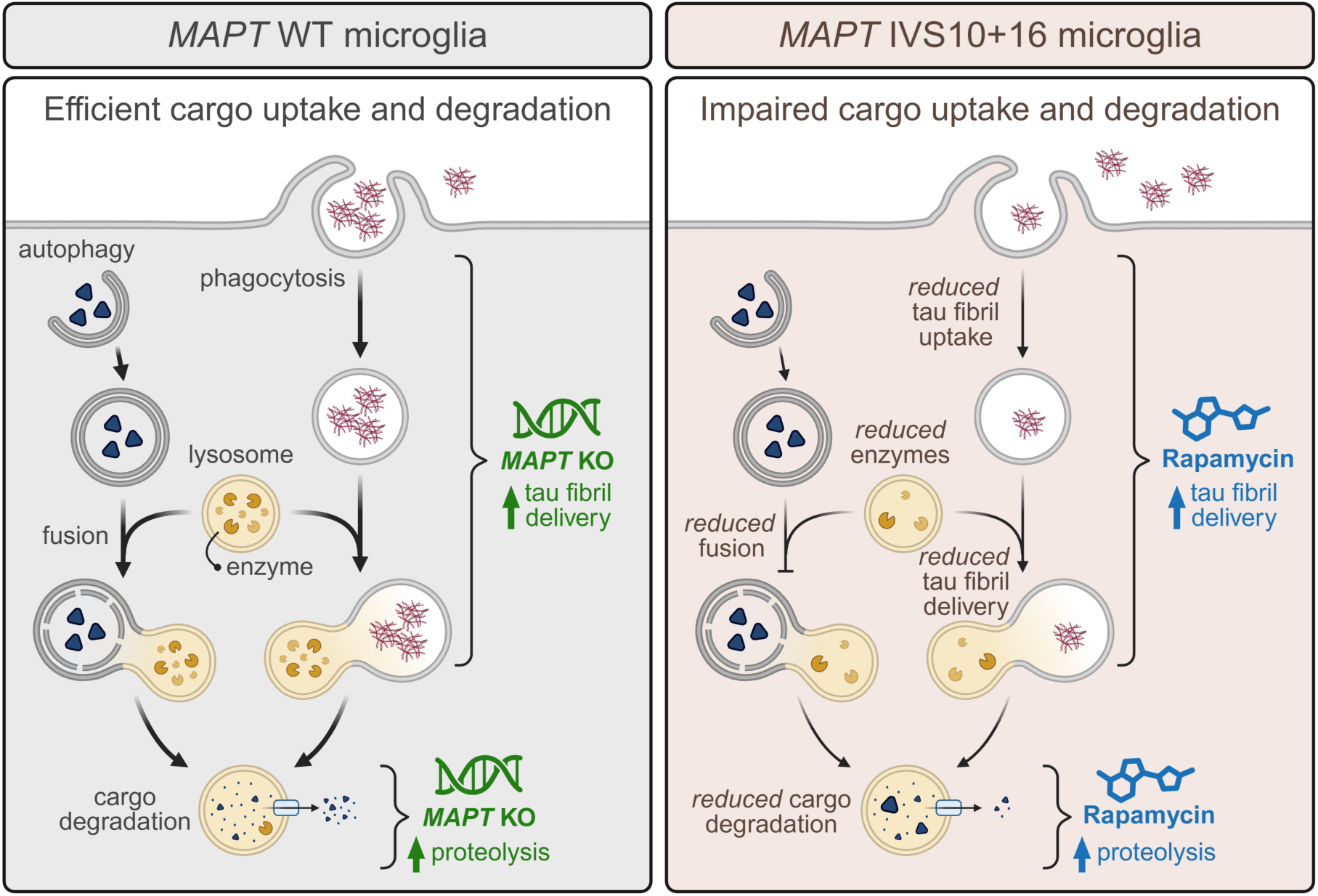
Model of *MAPT*-mediated regulation of lysosomal function and phagocytosis in microglia. *MAPT* IVS10+16 impairs lysosomal function and autophagy, while genetic *MAPT* loss enhances phagocytosis and degradative capacity.

As the resident macrophages of the CNS, microglia are responsible for surveilling the brain milieu to phagocytose and degrade waste in health and disease. As such, microglia are increasingly recognized as active regulators of tau pathophysiology through their capacity to internalize and process extracellular tau and tau-containing cellular material. Microglia activation is evident in FTLD-tau and has been detected in presymptomatic *MAPT* mutation carriers, suggesting that microglial responses emerge early in disease^60–65^. Microglia can remove extracellular tau and engulf tau-burdened neurons and synapses^11,66,67^, whereas inefficient processing of internalized tau may permit its persistence or release with consequences for tau propagation^14,68,69^. Although much of this work has examined microglial responses to neuronal-derived pathology, human microglia express the *MAPT* gene and tau protein, and *MAPT* IVS10+16 produces cell-autonomous abnormalities in phagocytosis, cytoskeletal organization, TREM2 signaling, and metabolism^45^. The present findings extend this work by identifying autophagy-endolysosomal degradation as an additional cell-autonomous process regulated by *MAPT* in microglia. This suggests that tau expressed within microglia may influence the fate of extracellular tau after uptake by determining the efficiency with which internalized cargo is delivered to and processed within degradative compartments.

The autophagy-endolysosomal network is increasingly implicated in tauopathy through genetic, proteomic, and functional studies^27–33,70–72^. Impairments in lysosomal and autophagy function have been reported in iPSC-derived neurons harboring *MAPT* mutations^73,74^, transdifferentiated neurons from Alzheimer’s disease patients^75^, and human primary neurons and astrocytes exposed to tau fibrils^76^. The present findings extend this relationship to human microglia and indicate that pathogenic *MAPT* alters degradative capacity without overtly reducing lysosome abundance or morphology. Instead, *MAPT* IVS10+16 suppressed lysosomal protease expression and activity, impaired autophagosome-lysosome fusion under autophagy-induced conditions, and limited processing of extracellular tau aggregates. This distinction between lysosome abundance and function is important because therapeutic approaches that depend on microglial uptake and degradation of proteopathic cargo may be constrained by the degradative state of the cell. Importantly, microglial lysosomal dysfunction was reversible. Rapamycin increased proteolytic activity and tau delivery to acidic compartments in mutant microglia, indicating that these functional deficits remain responsive to pathway modulation and that autophagy-lysosome function contributes to impaired tau processing. Together, these findings position microglia degradative capacity as a modifiable component of the cellular response to tau pathology. Future studies may explore the mechanistic relationship in microglia between lysosomal function and their other disease-modifying roles, such as exocytosis of pathological protein aggregates.

Interestingly, several defects became most apparent when the degradative system was challenged. While basal lysosome abundance, morphology, and autophagosome-lysosome colocalization were largely preserved, differences in lysosomal responses were accentuated following autophagic induction or tau fibril exposure. These findings suggest that *MAPT* IVS10+16 does not produce a generalized loss of lysosomes but instead limits the capacity of microglia to adapt their degradative machinery to increased cargo or proteopathic stress. Such loss of degradative reserve may be particularly relevant for microglia, which must rapidly increase phagocytic and lysosomal activity in response to cellular debris and protein aggregates.

Cathepsin proteases play critical roles in lysosomal degradation and contribute to CNS homeostasis, including the processing of aggregation-prone proteins such as tau. In *MAPT* IVS10+16 microglia, we observed reduced expression of lysosomal enzyme genes together with reduced cathepsin B and D protein levels. The importance of these proteases for CNS homeostasis is supported by human and animal studies: loss-of-function mutations in *CTSD* cause a rare, fatal neuronal ceroid lipofuscinosis disease^77,78^, while loss of cathepsins D or B in the CNS of mice is associated with microglia activation, accumulation of proteinopathy-related proteins including tau, and neuronal loss^56,79,80^. Conversely, excessive extracellular levels of cathepsins B and D secreted from microglia have neurotoxic effects^81,82^, indicating that cathepsin abundance and localization must be tightly regulated. Tau contains multiple cathepsin B cleavage sites and can be efficiently cleaved by cathepsin B *in vitro*, whereas cathepsin D has few predicted cleavage sites in tau and shows more limited tau proteolysis across a narrow pH range^83^. Thus, the reduction of both cathepsins in *MAPT* IVS10+16 microglia is more consistent with a broader impairment of lysosomal proteolytic capacity than with selective loss of enzymes that directly degrade tau.

Tau was initially defined by its role in microtubule binding and dynamics, but additional functions have since been described in neurons, including regulation of intracellular trafficking, neuronal excitability, nucleocytoplasmic transport, RNA metabolism, and genomic stability^84–100^. More recently, tau has also been shown to have cell-autonomous functions in glia, where endogenous tau supports lipid droplet formation, processing of peroxidated lipids, and resistance to oxidative stress through a microtubule-dependent role in lipid droplet budding from the endoplasmic reticulum^101^. Our findings extend the functional repertoire of tau to human microglia, where loss of *MAPT* increased lysosomal proteolytic activity and delivery of extracellular tau aggregates to acidic compartments. This effect may be cell-and cargo-dependent, as tau knockout has been reported to reduce autophagic clearance of Aβ in primary mouse hippocampal neurons^102^. Thus, the consequences of tau loss on degradative pathways likely depend on the cellular context, cargo, and the specific pathway engaged.

One possible mechanism by which *MAPT* mutations and tau loss could impact autophagy and tau uptake in microglia is altered microtubule-dependent trafficking. Autophagosome and lysosome transport and fusion rely on the microtubule network^103^, while phagocytosis requires coordinated cytoskeletal remodeling^104^. Consistent with this possibility, *MAPT* IVS10+16 microglia exhibit microtubule abnormalities^45^. However, tau also has microtubule-independent functions, and whether these contribute to regulation of lysosomal and phagocytic pathways in microglia remains to be determined.

The reciprocal phenotypes produced by *MAPT* IVS10+16 and *MAPT* knockout support a relationship between tau and microglial degradative state, but they should not be interpreted as opposite ends of a single linear pathway. *MAPT* IVS10+16 alters exon 10 splicing and increases 4R tau, whereas *MAPT* knockout removes endogenous tau. The mutant phenotype could therefore arise from altered tau isoform balance, changes in tau-microtubule interactions, gain of pathogenic function, or disruption of another physiological tau activity. Defining which of these mechanisms is responsible will be important for understanding how disease-causing *MAPT* variants alter microglial biology and whether similar mechanisms operate in sporadic tauopathies.

The reversibility of the lysosomal phenotype has potential therapeutic implications. Increasing autophagy-lysosome activity enhanced proteolytic function and tau processing in mutant microglia, while loss of *MAPT* increased the same degradative measures. These findings raise the possibility that lysosomal enhancement or tau lowering could improve selected microglial degradative functions. However, the broader consequences of reducing tau in glia require careful consideration. Endogenous tau supports lipid droplet formation and lipid handling in other glial populations^101^, indicating that tau can contribute to protective glial stress responses in a pathway-and cell-type-dependent manner. Thus, approaches that lower tau may enhance some aspects of microglial degradation while affecting other homeostatic functions. Defining the consequences of partial versus complete tau reduction across glial populations will therefore be important for therapeutic strategies targeting tau.

The extent to which these findings generalize beyond familial *MAPT* tauopathy remains unknown. *MAPT* IVS10+16 provides a powerful system in which altered tau biology is genetically defined and precedes neurodegeneration, but sporadic tauopathies arise through more heterogeneous mechanisms. Determining whether similar microglial degradative defects occur in PSP, CBD, Alzheimer’s disease, and other tauopathies will establish whether impaired lysosomal adaptation represents a broader feature of tau pathology.

Overall, this study used complementary *MAPT* IVS10+16 and *MAPT* knockout human microglia to define a bidirectional relationship between tau and lysosomal degradative function. These findings identify microglial lysosomal dysfunction as a cell-intrinsic consequence of pathogenic *MAPT* signaling that may limit the capacity of microglia to respond to and process proteopathic cargo. More broadly, they position microglial degradative pathways as modifiable determinants of tau handling and support lysosomal enhancement and tau-lowering strategies as potential therapeutic approaches for tauopathies.

## Methods

The research described in this study complies with all relevant ethical regulations and was approved by the Washington University School of Medicine (IRB 201104178 and 201306108).

### iPSC lines

Human iPSC lines were generated from dermal fibroblasts using non-integrating Sendai virus as previously described^46^. All iPSC lines used in this study underwent standard quality control, including assessment of pluripotency markers by immunocytochemistry (ICC), karyotyping to exclude chromosomal abnormalities, and confirmation of genotype by Sanger sequencing (**Supplemental Figure 1**)^45,46^.

To study the impact of *MAPT* IVS10+16 on protein degradation pathways, we utilized three independent donor iPSC backgrounds and their corresponding isogenic pairs: GIH36C2, GIH178C1, and KOLF2.1J (**Supplemental Table 1**). Across these backgrounds, isogenic pairs were generated using complementary strategies to model the *MAPT* IVS10+16 mutation. Two pairs were derived from patient lines carrying one copy of the *MAPT* IVS10+16 mutation (GIH36C2 and GIH178C1) and were subsequently corrected to WT using CRIPSR/Cas9 (GIH36C2d1D01 and GIH178C1d2A10, respectively). A third pair was generated by engineering one copy of the *MAPT* IVS10+16 mutation (KOLF2.1J.IVS10+16d2F04) into the control KOLF2.1J background (JAX JIPSC1000^47^), providing an orthogonal approach to assess mutation effects independent of patient specific background variation.

To examine the effects of *MAPT* loss on protein degradation pathways, we utilized a *MAPT* knockout (KO) model generated on the control KOLF2.1J WT background (**Supplemental Table 1**). The *MAPT* KO (JAX JIPSC001670) was engineered in the KOLF2.1J background using Cas9-mediated deletion with two ‘PAM-out’ guide RNAs flanking the *MAPT* locus and a bridging repair oligonucleotide to excise the majority of the coding sequence. The bridging oligo incorporates degenerate bases at the deletion junction, enabling identification of homozygous KO clones carrying two copies of the engineered deletion by Sanger sequencing.

iPSCs were maintained in mTesR1 medium (STEMCell Technologies 85850) on Cultrex PathClear Basement Membrane Extract (R&D Systems 3532-010-02) and passaged with Accutase (Innovative Cell Technologies AT-104). Karyotyping and Sanger sequencing to verify mutation status were performed every 10 passages, and cultures were confirmed to be free of mycoplasma with monthly testing.

### Generation of iPSC-derived microglia (iMG)

Human iPSCs were differentiated into iMG using a two-step protocol^48^. In the first step, iPSCs were reprogrammed into HPC using a STEMdiff Hematopoietic kit (STEMCell Technologies 05310) following the manufacturer’s instructions as previously reported^45,48^. Briefly, confluent iPSCs were detached with ReLeSR (STEMCell Technologies 100-0484) and plated at 40 aggregates per well in 6-well plates in mTeSR1 supplemented with 10uM Rock inhibitor Y-27632 dihydrochloride (STEMCell Technologies 72302) on Day -1. On Day 0, cells were transferred to Medium A (STEMdiff Hematopoietic kit) to induce mesodermal patterning. On Day 3, cultures were transferred to Medium B to promote HPC differentiation, with full media replacement performed again on Day 5. From Days 7-12, fresh Medium B was added every two days without media removal to support expansion of floating HPCs. On Day 12, the floating cell HPC population was collected and cryopreserved in CyroStor CS10 (BioLife Solutions 210374). From each differentiation, 100,000 cells were reserved for validation of HPC markers by flow cytometry, and 50,000 cells were reserved for Sanger sequencing to confirm mutation status.

HPCs were differentiated into iMG as previously reported, with slight modifications^45,48^. HPCs were thawed and plated at 150,000-350,000 cells per well (number optimized for each iPSC donor) in Cultrex-coated 6-well plates in microglia basal medium (DMEM/F12, Sigma-Aldrich D8437; 2x insulin-transferrin-selenite solution, Thermo Fisher 41400045; 1x B27 Supplement, Thermo Fisher 17504044; 1x N2 Supplement, Thermo Fisher 17502001; 1x Glutamax, Thermo Fisher 35050061; 1x non-essential amino acids, Thermo Fisher 11140050; 1x Penicillin-Streptomycin, Thermo Fisher 15140122; 400uM monothioglycerol, Sigma Aldrich M1753; 5ug/mL recombinant human insulin, Sigma Aldrich I2643) supplemented with maintenance cytokines (100ng/mL IL-34, PeproTech 200-34; 50ng/mL TGF-β1, PeproTech 100-21; 25ng/mL M-CSF, PeproTech 300-25). The day of HPC thaw was designated Day 12.

Cells were then maintained in microglia basal medium supplemented with fresh maintenance cytokines from Days 12-37, with fresh media added every two days. Cultures were expanded on Days 18, 24, and 30 by evenly splitting both floating and adherent populations based on cell density. On Day 37, cells were transitioned to maturation conditions with microglia basal medium supplemented with maintenance cytokines and maturation factors (100ng/mL CD200, Novoprotein C311; 100ng/mL CX3CL1, PeproTech 300-31).

On Day 39, iMG were replated at equal densities for experiments. To replate iMG, the floating cell population was collected and combined with adherent cells that were lifted with DPBS (Sigma Aldrich D8537). Combined pools were centrifuged at 300g for 5min at RT and resuspended in fresh maturation media. Cell counts were determined using Invitrogen Countess 3 automated cell counter. From Days 12-39, cultures exhibiting signs of stress (i.e. debris, rounding) were fed with 2x concentration of maintenance cytokines; this was routinely required for iMG derived from the KOLF2.1J background. All experiments were performed in mature iMG on Days 40-41.

### Evaluation of HPC and iMG cell surface markers

HPCs and iMG were analyzed for lineage-specific surface markers by flow cytometry at Day 12 and Day 40, respectively (**Supplemental Figure 2A-B**). Cells were stained with Zombie Aqua viability dye (1:1000, BioLegend 423102) diluted in DPBS for 15min at 4°C in the dark. Cells were washed with additional DPBS and centrifuged at 500g for 3min at 4°C (HPCs) or 300g for 5min at 4°C (iMG). Cells were then incubated with Fc receptor blocking reagent (1:1000, Milteny Biotec 130-059-901) diluted in FACS buffer (2% FBS [Biowest S1620] in DPBS, sterile filtered) for 10min at 4°C. After washing with FACS buffer, HPCs were stained with CD34-FITC (1:200, BioLegend 343504), CD43-APC (1:200, BioLegend 343206), and CD45-Alexa Fluor 700 (1:200, BioLegend 304024) in FACS buffer for 20min at 4°C. iMG were stained with CD11b-PE/Cy7 (1:300, BioLegend 101216) and CD45-Alexa Fluor 700 (1:300) in FACS buffer for 20min at 4°C. Cells were washed and resuspended in fresh FACS buffer for acquisition. Cell surface markers were acquired on a FACS Symphony A1 analyzer. Single color-stained beads and unstained control cells were used for compensation. The resulting data were analyzed with FlowJo software (v10).

### Real Time-PCR for *MAPT* expression

*MAPT* expression in iMG derived from the *MAPT* KO iPSC line was assessed via Real Time-PCR, with *MAPT* WT iMG and iPSC-derived neuron samples included as positive controls. cDNA was generated from RNA using a High-Capacity cDNA Reverse Transcription Kit (Thermo Fisher 4368813) and then amplified using primers targeting *MAPT* and *GAPDH* as a control: *MAPT* forward, 5’-AAGTCGCCGTCTTCCGCCAAG-3’; *MAPT* reverse, 5’-GTCCAGGGACCCAATCTTCGA-3’; *GAPDH* forward, 5’-ATGTTCGTCATGGGTGTGAA-3’; *GAPDH* reverse, 5’-TGTGGTCATGAGTCCTTCCA-3’^105^. The PCR amplification was performed using 200ng cDNA for iMG samples and 100ng cDNA for iNeuron samples per reaction and a GoTaq Flexi DNA Polymerase Kit (Promega M8296) in a final volume of 25uL. PCR products were resolved on a 1.5% agarose gel for 60min.

### Tau preformed fibrils (tau PFFs)

Tau uptake and trafficking to acidic vesicles were assessed using fluorescently labeled tau pre-formed fibrils (PFFs). Tau PFFs labeled with ATTO488 were used to quantify cellular tau uptake. Tau PFF-ATTO488 consisted of human recombinant Tau-441 (2N4R, P301S) fibrils produced in *E. coli* and fluorescently labeled (StressMarq Biosciences, SPR-329-A488).

Tau PFFs labeled with pHrodo were used to assess delivery to acidic vesicles. Tau PFF-pHrodo were generated from human recombinant Tau-441 (2N4R, P301S) fibrils produced in baculovirus/Sf9 cells (StressMarq Biosciences, SPR-471) by conjugation to pHrodo Red succinimidyl ester (Thermo Fisher Scientific, P36011). Briefly, 1 mg/mL tau PFFs in 25 mM sodium bicarbonate buffer were incubated with a 5-fold molar excess of pHrodo dye overnight at room temperature. Excess dye was removed by dialysis using Slide-A-Lyzer cassettes (3.5 kDa MWCO; Thermo Fisher Scientific, 66335) according to the manufacturer’s instructions.

Prior to treatment, ATTO488-and pHrodo-labeled PFFs were sonicated (QSonica bath sonicator; 20% amplitude, 30s on/off ×3 cycles) to generate fibrillar species suitable for cellular uptake.

### RNA sequencing

On Day 40 of microglia differentiation, iMG were treated with 50nM tau PFF-ATTO488 or with equivalent volume DPBS as a vehicle control for 24hrs. Following treatment, cells were washed and then collected in Qiazol lysis reagent (Qiagen 79306). RNA extraction was performed using RNeasy Mini Kit (Qiagen 74104), and the total RNA integrity was measured using Agilent Bioanalyzer or 4200 Tapestation. Library preparation was performed using ribosome depletion methods (Ribo-ZERO, Illumina-EpiCentre) and sequenced on an Illumina NovaSeq-6000 using paired end reads extending 150 bases. Three samples per group, representing independent wells from a single differentiation of the GIH36C2 donor, were submitted for sequencing. One vehicle-treated WT sample failed quality control and was excluded prior to principal component and differential gene expression analyses.

### Principal component, differential expression, and pathway analyses

Transcriptome analyses were based on 15,499 protein-coding genes (≥1 CPM in > 50% sequencing cohort). PCAs were calculated using the top 500 most variable genes across all groups using rlog-normalized counts (DESeq2^106^). Differential gene expression was performed using DESeq2 for R (v4.3.1) for the following comparisons: *MAPT* IVS10+16 Veh vs WT Veh, WT Tau PFF vs WT Veh, IVS10+16 Tau PFF vs IVS10+16 Veh, and IVS10+16 Tau PFF vs WT Tau PFF. PCA and volcano plots were created using ggplot2^107^. Genes passing the Benjamini-Hochberg adjusted p-value FDR≤0.05 threshold were analyzed using Enrichr against KEGG (2021 Human) and GO biological processes datasets^108^.

### Immunoblotting

To evaluate the impact of *MAPT* IVS10+16 on protein expression, iMG were plated at 500,000 cells per well in 6-well plates on Day 39 and harvested on Day 41 by pooling three wells (1.5 × 10^6^ cells per line per differentiation). Cell pellets were snap frozen at -80°C. Cells were lysed in RIPA buffer (Teknova R3792) supplemented with PhoSTOP phosphatase inhibitor (1x; Roche 4906845001), N-ethylmaleimide (0.625 mg/mL; Fisher Scientific AC156100050), and protease inhibitor (1:500; Sigma P8340). Lysates were sonicated (QSonica bath sonicator; 50% amplitude, 30s on/off ×4 cycles) and clarified by centrifugation (14,000g, 15 min, 4°C). Total protein concentrations were measured using a BCA assay (Thermo Fisher 23225).

Equal amounts of protein (20μg) were mixed with 4x Bolt LDS sample buffer (Invitrogen B0007) containing 10% β-mercaptoethanol (Sigma M-7154) and heated at 70°C for 10min. Samples were resolved on 4-12% Bis-Tris Plus gels (Invitrogen NW04120BOX) using MES SDS running buffer (Invitrogen NP0002) and a dual color protein standard (Invitrogen LC5602) at 100V for 1hr 30min. Proteins were then transferred to 0.2μm PVDF membranes (Millipore ISEQ00010) using NuPAGE transfer buffer (Thermo Fisher NP0006) with 20% methanol (Fisher Chemical A412P-4). Membranes were blocked in 3% BSA (Sigma Aldrich A2153-100G) in PBS-T (0.1% Tween-20; Sigma P1379) and incubated with primary antibodies overnight at 4°C. After washing, membranes were incubated with HRP-conjugated secondary antibodies (1:3000) for 1hr at RT, washed, and developed using SuperSignal West Pico PLUS (Thermo Fisher 3458) or Lumigen ECL Ultra (Fisher Scientific NC0240697). Membranes were stripped using Restore PLUS buffer (Thermo Scientific 46430) prior to reprobing.

The following primary antibodies were used: LAMP1 D2D11 (rabbit, 1:2000, Cell Signaling Technology 9091S), LAMP2A AMC2 (rabbit, 1:3000, Thermo Fisher 51-2200), Cathepsin D (rabbit, 1:1000, Cell Signaling Technologies 2284), Cathepsin B D1C7Y (rabbit, 1:1000, Cell Signaling Technologies 31718), GAPDH (mouse, 1:1000, Invitrogen MA5-15738). The following secondary antibodies were used: Goat anti-mouse IgG (H+L)-HRP conjugate (1:3000, BioRad 1706516), goat anti-rabbit IgG (H+L)-HRP conjugate (1:3000, BioRad 1706515), horse anti-mouse IgG HRP-linked antibody (1:3000, Cell Signaling Technology 7076), goat anti-rabbit IgG HRP-linked antibody (1:3000, Cell Signaling Technology 7074).

### Immunocytochemistry

To evaluate protein localization and subcellular organization in iMG, immunocytochemistry was performed. iMG were plated at 80,000 cells per well on glass coverslips in 24-well plates on Day 39. On Day 40, cells were fixed by adding 16% paraformaldehyde (PFA, Electron Microscopy Sciences 15710) directly into the culture media for a final concentration of 4% and incubated for 20min at RT. Cells were then washed with DPBS and permeabilized with 0.5% Triton-X100 (Sigma Aldrich T8787) in DPBS for 20min. Cells were blocked in blocking buffer (3% BSA and 0.3% Triton-X100 in DPBS) for 1hr at RT. Cells were incubated with primary antibodies diluted in blocking buffer overnight at 4°C in the dark. The following day, cells were washed with DPBS and incubated with secondary antibodies diluted in blocking buffer for 1hr at RT in the dark. After washing, nuclei were stained with NucBlue Fixed Cell ReadyProbes Reagent (Invitrogen R37606) solution (2 drops/mL in DPBS) for 15min at RT, followed by a final DPBS wash. Coverslips were then mounted onto slides using Fluoromount-G (Invitrogen 00-4958-02) and stored at 4°C prior to imaging.

Confocal imaging was performed on a Nikon AX-R with NSPARC using the NSPARC detector in SR mode, 8k Galvano scanner, and 60x oil-immersion objective. Images (1024×1024 pixels) were acquired at 4.6x zoom, 1 integration, 0.4ms dwell time. Z-stack images were collected across the full cell volume (0.15um step size). Channels were acquired sequentially. Following acquisition, images were deconvolved in NIS Elements using blind 3D deconvolution over 20 iterations. Within an experiment, all samples were imaged using identical acquisition parameters during a single imaging session for direct comparison.

The following primary antibodies were used: IBA1 (goat, 1:500, Abcam 5076), LAMP1 D2D11 (rabbit, 1:200, Cell Signaling Technology 9091S), LAMP1 (mouse, 1:100, Cell Signaling Technology 15665S), LC3B (rabbit, 1:400, Cell Signaling Technology 2775), TMEM119 (rabbit, 1:100, Abcam ab185333). The following secondary antibodies were used: Donkey anti-Mouse IgG Alexa Fluor 488 (Invitrogen A21202), donkey anti-Rabbit IgG Alexa Fluor 647 (Invitrogen A31573), donkey anti-Goat IgG Alexa Fluor 555 (Invitrogen A21432).

### Quantification of lysosomal morphometry

LAMP1 intensity, vesicle size and vesicle number were quantified using Imaris (v10.2). A 3D Cell module was applied in batch to all images within each experiment, defining the cytoplasm (IBA1; Surface module), nucleus (NucBlue, Surface module), and LAMP1 vesicles (Spot module). Total LAMP1 intensity was measured per cell and normalized to cell volume. Vesicle number and mean volume per cell were also quantified.

### Monitoring autophagic vesicles

Autophagic vesicles were assessed using a CYTO-ID Autophagy Detection kit (Enzo ENZ-51031). For flow cytometry, iMG were plated at 150,000 cells per well in 24-well plates. For live imaging, iMG were plated at 200,000 cells per dish in 35 mm dishes with 20mm #1.5 glass coverslip bottoms. iMG were treated with rapamycin (500nM) and chloroquine (10uM) or an equivalent volume of vehicle (DMSO) for 6hrs. CYTO-ID dye (1:1000) was applied for 20min at 37°C.

To evaluate CYTO-ID by flow cytometry, cells were collected on ice, washed with DPBS, and stained with Zombie Aqua dye (1:1000 in DPBS) for 20 min prior to resuspension in FACS buffer. Samples, along with single color and unstained controls, were acquired on a FACS Symphony A1 analyzer and analyzed using FlowJo software (v10). CYTO-ID positivity was confirmed to be 100% in the live cell (Zombie Aqua-) population across biological replicates. CYTO-ID was quantified as MFI within the live cell population.

To evaluate the number and size of autophagic vesicles, CYTO-ID was assessed by live cell confocal imaging. Cells were co-labelled with SPY555-actin (Cytoskeleton Inc. CY-SC202, 1:1000, 2hrs) and Hoechst (33342, 1:2000, 20min) to visualize the actin cytoskeleton and nuclei, respectively. Treatments and labeling were staggered across replicates to ensure consistent incubation times, and media were replaced immediately prior to imaging. Confocal imaging was performed on a Nikon AX-R microscope with NSPARC using the NSPARC detector in SR mode, a 2k resonant scanner, and a 60x oil-immersion objective (2.5x zoom; 2048×2048 pixels). Channels were acquired simultaneously in single z-slice images. Within an experiment, all samples were imaged using identical acquisition parameters during a single imaging session. Image analyses were performed in Imaris (v10.2). CYTO-ID+ vesicles were detected using a Spot module, and nuclei were detected using a Surface module, with parameters applied in batch across all images. Vesicle counts were normalized to nuclei per field, and mean vesicle area was calculated per field.

### Autophagosome-lysosome fusion

To assess autophagosome-lysosome fusion, we quantified co-localization of LC3B-positive autophagosomes with LAMP1-positive lysosomes in iMG. Rapamycin (Enzo 51031-RAP-25) was used to stimulate autophagic flux, and bafilomycin A1 (Fisher Scientific 50-186-9239) to inhibit lysosomal acidification, providing a functional control to define assay dynamic range and interpret changes in co-localization. Rapamycin (1uM) and bafilomycin (400nM) or vehicle (DMSO) were added to iMG for 4hrs prior to fixation. Co-localization was quantified in Imaris (v10.2). LAMP1 and LC3B were first segmented using Surface modules. The Surface-Surface Co-localization XTension was then applied to generate a co-localization surface representing overlapping voxels. The volume of the co-localization surface was normalized to the LAMP1 surface volume within each field, yielding the fraction of LAMP1 which co-localizes with LC3B.

### Assessment of lysosomes following tau PFF exposure

To determine how tau PFFs impact lysosomes, iMG were plated at 100,000 cells per well in 48-well plates on Day 39. On Day 40, iMG were treated with 500nM tau PFF-ATTO488 or an equivalent volume vehicle (DPBS) for 24hrs. During the final 30min of treatment, cells were labeled with 100nM LysoTracker Red (Thermo Fisher L7528). Cells were then collected on ice, washed with DPBS, and stained with Zombie Aqua dye (1:1000 in DPBS) for 20min prior to resuspension in FACS buffer for acquisition. Samples, along with single color and unstained controls, were acquired on a FACS Symphony A1 analyzer and analyzed with FlowJo software (v10). The percentage of tau PFF+ cells and the tau PFF geometric mean fluorescence intensity (MFI) were quantified within the live cell (Zombie Aqua-) population. LysoTracker positivity was confirmed to be 100% across live cells for each biological replicate. The LysoTracker MFI was quantified within live cells for vehicle-treated samples and within live, tau PFF+ cells for tau-treated samples.

### Live-cell imaging of proteolytic activity and tau uptake

For Incucyte live imaging, iMG were plated at 50,000 cells per well in 96-well plates. For rapamycin rescue experiments, iMG were pre-treated with 100nM rapamycin or equivalent volume vehicle (DMSO) 4hrs prior to substrate addition. Fluorescent substrates were added to the culture medium directly before imaging on an Incucyte S3, acquiring 4-5 images per well every hour for 12-24hrs. Proteolytic capacity was assessed using DQ Red BSA (1:1000 to produce 1ug/mL; Thermo Fisher D12051). Cathepsin B activity was measured using Magic Red Fluorescent Cathepsin B Assay Kit (1:1000; Antibodies Inc 938). Tau fibril delivery to acidic vesicles was evaluated using tau PFF-pHrodo (250nM). Quantification was performed using Incucyte software (v2025B). iMG were segmented from phase images and fluorescent signal from substrate channels using Surface Fit segmentation and thresholding. Total integrated fluorescence intensity was normalized to cell area.

## Statistics and reproducibility

Data visualization and statistical testing was performed in GraphPad Prism (v11). Data are presented as individual values and mean ± SEM. When comparing two groups, two-tailed paired or unpaired t-tests were used. Equal variance was not assumed. When the F-test to compare variances yielded p≤0.05, a two-tailed Welch’s t-test was applied. When comparing multiple groups across two factors (e.g. genotype and treatment), a two-way ANOVA was performed followed by Tukey’s multiple comparisons test or by Fisher’s LSD test when a subset of post-hoc comparisons was performed. The data distribution was assumed to be normal. Specific statistical tests are detailed in the Figure and Supplemental Figure legends.

## Data Availability

All data generated or analyzed during this study are included in this published article (and its supplementary information files).

## Acknowledgments

We would like to thank the research subjects and their families who generously participated in this study. We thank Torri Ball for administrative support and careful review of the study. We thank Dr. Samantha Swift for thoughtful discussions throughout the study. This work was supported by access to equipment made possible by the Hope Center for Neurological Disorders and the Departments of Neurology and Psychiatry at Washington University School of Medicine. Confocal images were generated on a Nikon AXR with NSPARC through the use of Washington University Center for Cellular Imaging-Neuro supported by Washington University School of Medicine, the Children’s Discovery Institute of Washington University and St. Louis Children’s Hospital (CDI-CORE-2015-505 and CDI-CORE-2019-813), and the Foundation for Barnes-Jewish Hospital (3770 and 4642). The content of this publication is solely the responsibility of the authors and does not necessarily represent the official views of the NIH. Funding provided by the National Institutes of Health (NS123985 [CMK], NS110890 [CMK], AG066444 [CMK], TR002345), the Knight ADRC Developmental Project in collaboration with the McDonnell Center for Cellular & Molecular Neurobiology [AKI], CurePSP [AKI], and the Rainwater Charitable Organization. Diagrams were generated using BioRender.com.

## Author Contributions

Designed experiments: KJS, AKI, CMK. Performed and analyzed experiments: KJS, RZ, AES, ES, AKI. Provided experimental tools: ST. Provided funding: CMK, AKI. Wrote the manuscript: KJS, CMK. Revised and approved manuscript: KJS, RZ, AES, ES, JAM, DJK, ST, AKI, CMK.

## Competing Interests

C.M.K. serves as an advisor for Eisai Co. Ltd and Synapticure Inc. The following authors have no competing interests: KJS, RZ, AES, ES, JAM, ST, DJK and AKI.

## Notes

### Competing Interest Statement

CMK serves as an advisor for Eisai Co. Ltd and Synapticure Inc. The following authors have no competing interests: KJS, RZ, AES, ES, JAM, ST, DJK and AKI.

