## Supplemental Figures for "*MAPT* regulates autophagic-lysosomal function and phagocytosis in human microglia"

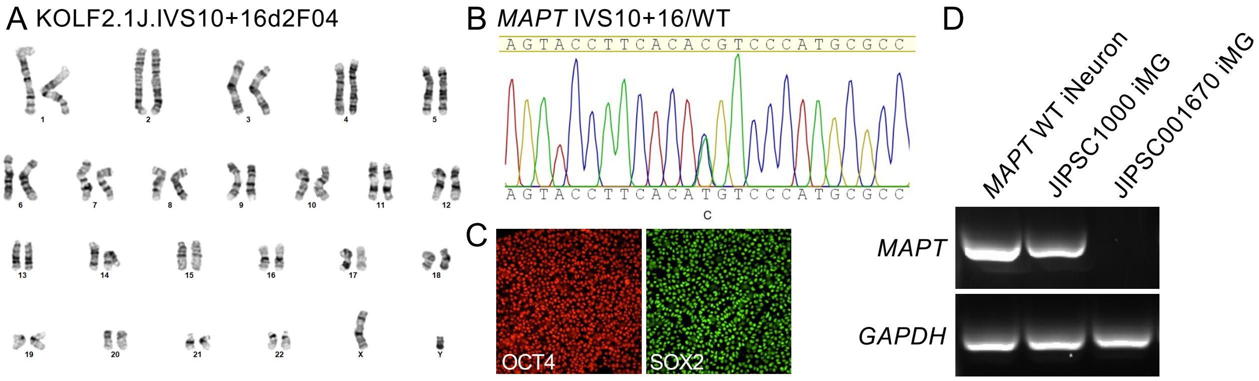


**Supplemental Figure 1. Characterization of newly generated iPSC lines**. A-C. Newly generated engineered iPSC line for studying the impact of *MAPT* IVS10+16 on lysosomal phenotypes. iPSC KOLF2.1J.IVS10+16d2F04 was engineered from *MAPT* WT/WT to *MAPT* IVS10+16/WT. A. G-band karyotyping confirmed chromosomal stability. B. Sanger sequencing at the *MAPT* locus confirmed a heterozygous edit to IVS10+16 in the *MAPT* locus. C. Immunocytochemistry confirmed expression of pluripotency markers. Additional iPSC lines used in this study have been characterized and reported previously: patient derived *MAPT* IVS10+16/WT (GIH36C2 and GIH178C1[45,46]); CRISPR/Cas9 corrected *MAPT* WT/WT (GIH36C2d1D01 and GIH178C1d2A10, respectively[45,46]); control *MAPT* WT/WT (KOLF2.1J; JAX Cat# JIPSC1000[47]); and MAPT KO (JAX Cat# JIPSC001670). D. RT-PCR using primers for *MAPT* and *GAPDH* confirmed loss of *MAPT* expression in JIPSC001670 iMG. *MAPT* WT iNeuron and isogenic *MAPT* WT JIPSC1000 iMG were included as positive controls for *MAPT* expression.

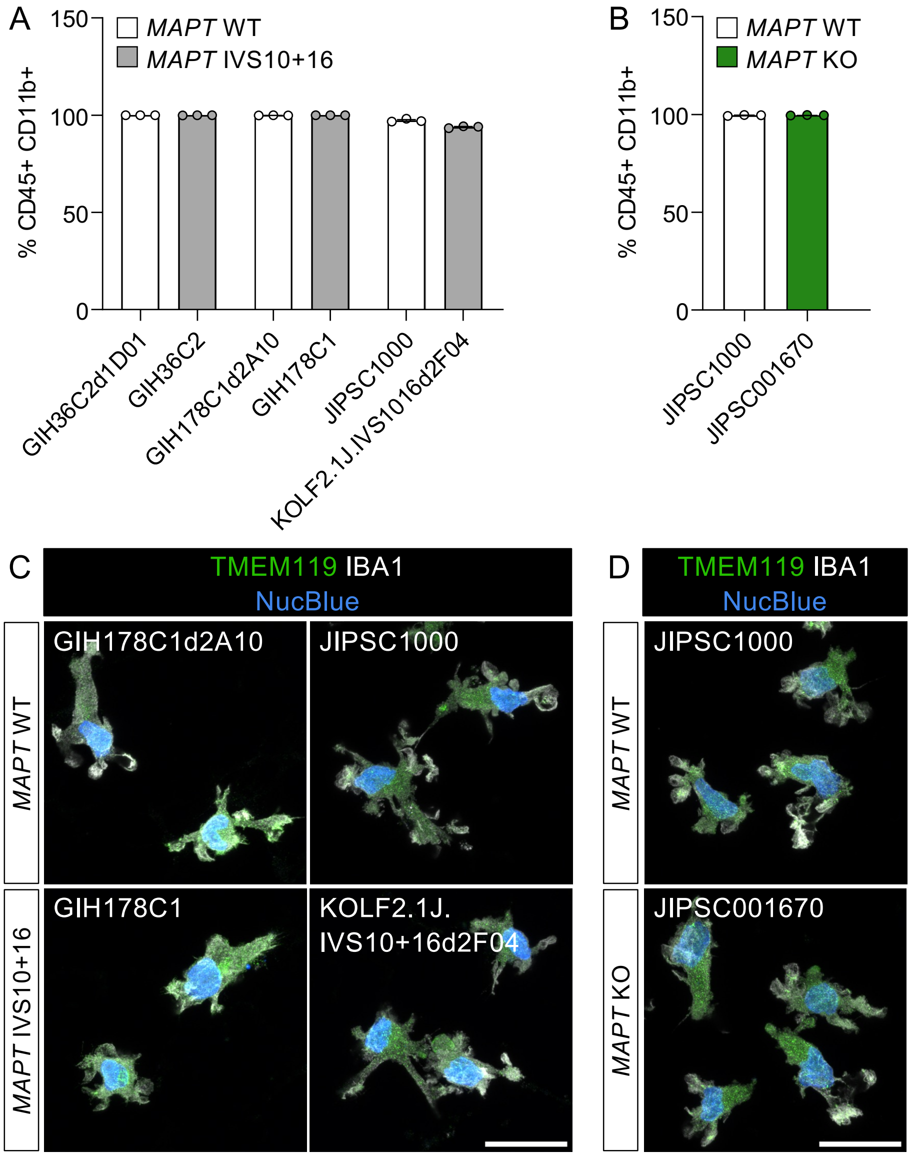


**Supplemental Figure 2. iMG differentiation validation across iPSC donors**. A-B. Each iPSC line used in the study was verified to differentiate into CD45 and CD11b double positive iMG by flow cytometry. A. *MAPT* IVS10+16 lines and isogenic controls. B. *MAPT* KO and isogenic control. C-D. Each iPSC line was verified to differentiate into TMEM119 (green) and IBA1 (white) double-positive iMG by immunocytochemistry. Representative max z projections with nuclei (NucBlue, blue). Scale bar, 20µm. C. *MAPT* IVS10+16 lines and isogenic controls. D. *MAPT* KO and isogenic control.

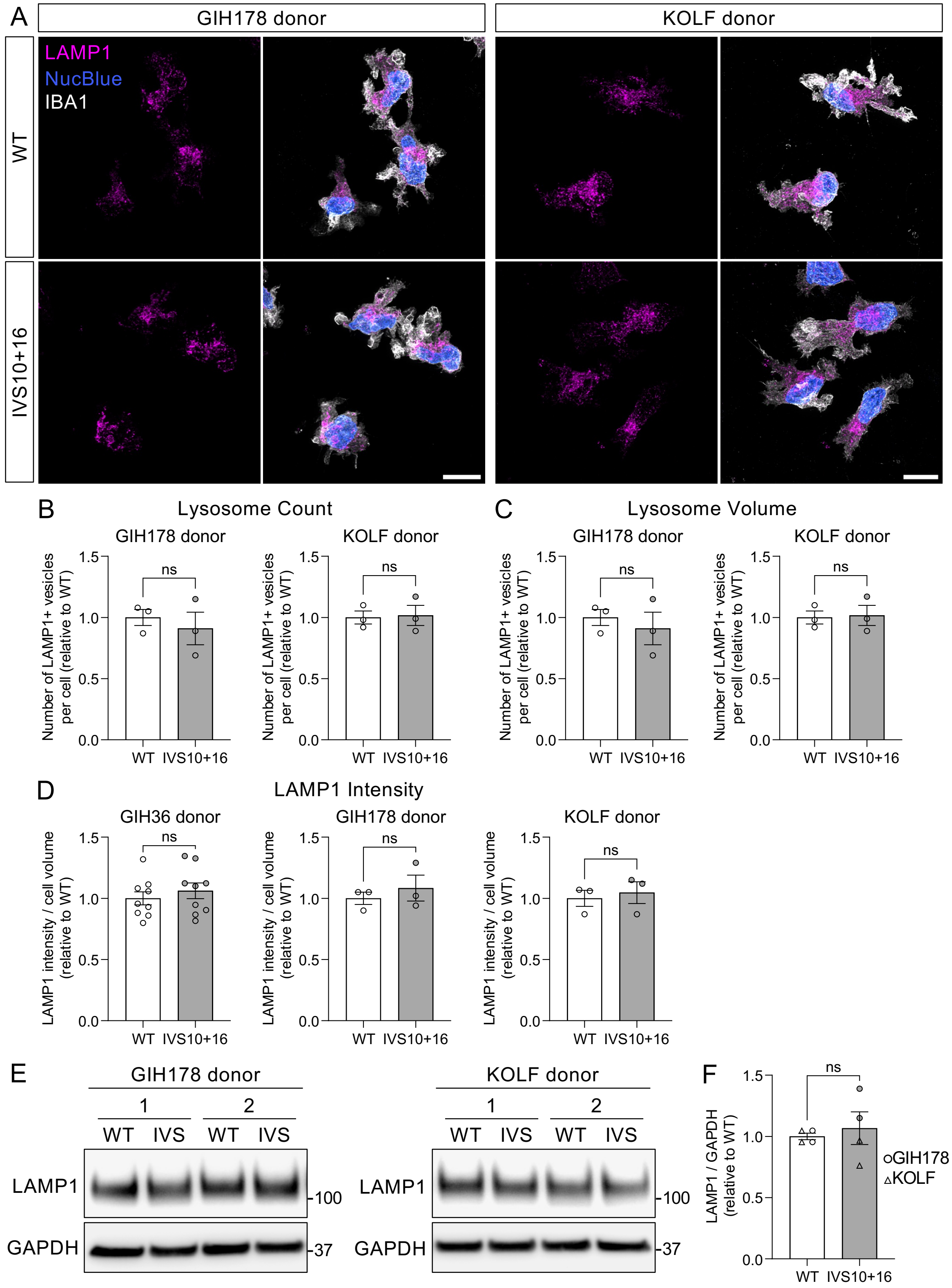
A
GIH178 donor
KOLF donor
LAMP1
NucBlue
IBA1
WT
IVS10+16
B
Lysosome Count
GIH178 donor
KOLF donor
Number of LAMP1+ vesicles per cell (relative to WT)
ns
WT
IVS10+16
C
Lysosome Volume
D
LAMP1 Intensity
GIH36 donor
LAMP1 intensity / cell volume (relative to WT)
E
1
2
WT
IVS
LAMP1
GAPDH
100
37
F
LAMP1 / GAPDH (relative to WT)
GIH178
KOLF

**Supplemental Figure 3. Lysosomal abundance and morphology are preserved in *MAPT* IVS10+16 microglia in independent iPSC donors (corresponding to Figure 2).** A-D. Immunocytochemistry for LAMP1-positive vesicles. A. Representative max z projections of LAMP1 (magenta), IBA1 (white), and nuclei (blue). Large field scale bar, 10µm. B. Quantification of LAMP1+ vesicle number per cell was performed in Imaris. Two-tailed unpaired t-tests; GIH178, p=0.5828; KOLF, p=0.8640. C. Quantification of LAMP1+ vesicle volume was performed in Imaris. Two-tailed unpaired t-tests; GIH178, p=0.6385; KOLF, p=0.1130. D. Quantification of LAMP1 sum intensity normalized to cell volume was performed in Imaris. Two-tailed unpaired t-tests; GIH36, p=0.4724; GIH178, p=0.5189; KOLF, p=0.6790. B-D. Data normalized to WT within each differentiation. GIH36: Data from three independent differentiations; each point represents an average from 3 wells. WT, n=269 cells; IVS10+16, n=268. GIH178: Data from one differentiation; each point represents one well. WT, n=43 cells; IVS10+16, n=34. KOLF: Data from one differentiation; each point represents one well. WT, n=35 cells; IVS10+16, n=33. E-F. Whole cell lysates were analyzed by SDS-PAGE immunoblotting. E. Representative immunoblots for LAMP1 (lysosomal membrane marker) and GAPDH (loading control) showing two independent differentiations per donor. F. LAMP1 protein levels quantified by densitometry, normalized to GAPDH, and expressed relative to WT controls. Data from four independent differentiations. GIH178, circles; KOLF, triangles. Two-tailed paired t-test (paired by donor and differentiation); p=0.6253. Graphs represent mean ± SEM. ns, not significant.

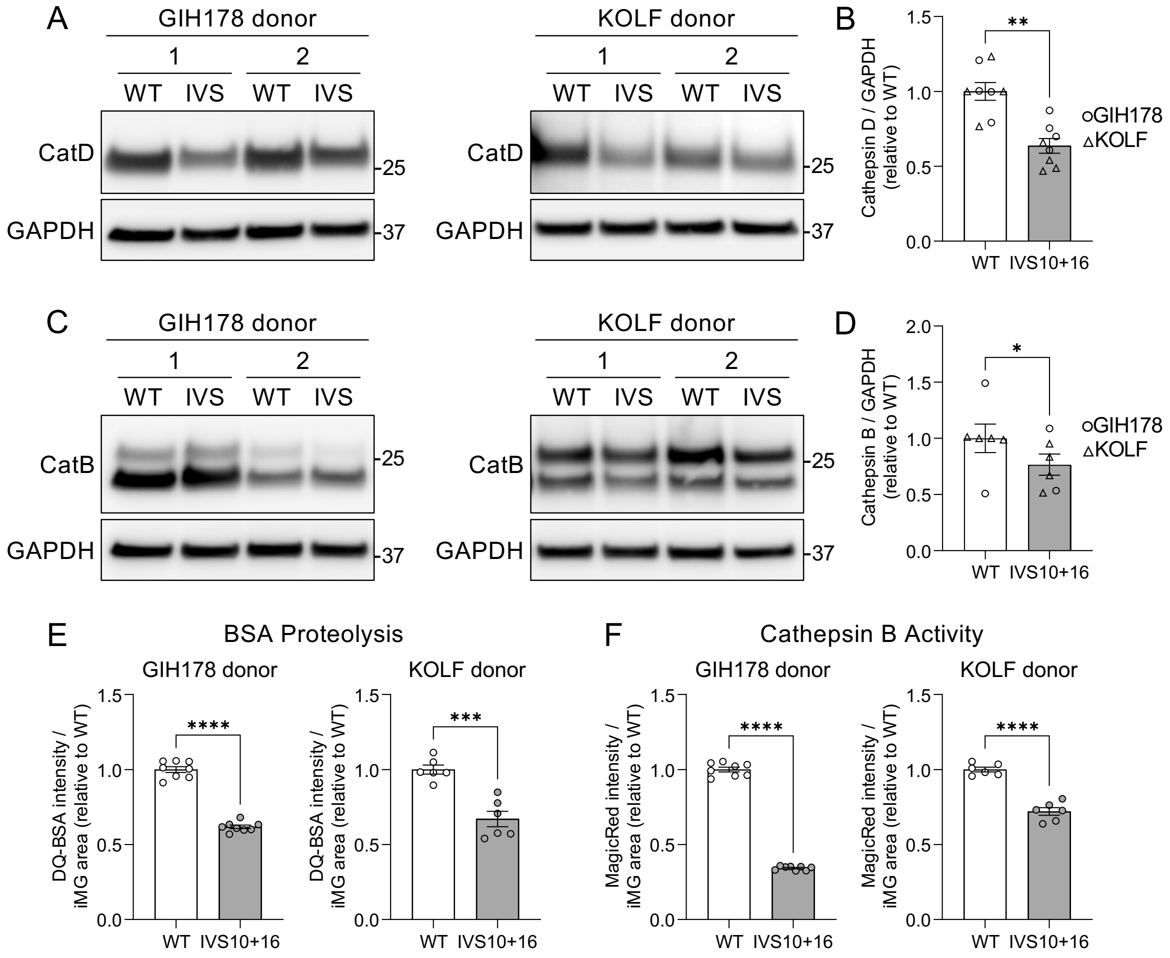


**Supplemental Figure 4. *MAPT* IVS10+16 disrupts lysosomal protease machinery and impairs degradative function in independent iPSC donors (corresponding to Figure 3).** A-D. Whole cell lysates were analyzed for cathepsin proteins by SDS-PAGE immunoblotting. GIH178, circles; KOLF, triangles. A. Representative immunoblot of cathepsin D showing two independent differentiations per donor. B. Cathepsin D protein levels quantified by densitometry, normalized to GAPDH, and expressed relative to WT. Data from eight independent differentiations. Two-tailed paired t-test (paired by donor and differentiation); p=0.0015. C. Representative immunoblot of cathepsin B showing two independent differentiations per donor. D. Cathepsin B protein levels quantified by densitometry, normalized to GAPDH, and expressed relative to WT. Data from six independent differentiations. Two-tailed paired t-test (paired by differentiation and donor); p=0.0367. E. iMG were treated with DQ-BSA (1µg/mL) and analyzed by Incucyte live cell imaging. Quantification of DQ-BSA integrated intensity at 24hrs was normalized to cell area and expressed relative to WT. Two-tailed unpaired t-tests; GIH178, p<0.0001, n=8 wells per genotype; KOLF2.1J, p=0.0003, n=6 wells per genotype. F. iMG were treated with Magic Red (1:1000) and analyzed by Incucyte live cell imaging. Quantification of Magic Red integrated intensity at 12hrs was normalized to cell area and expressed relative to WT. GIH178: two-tailed unpaired t-test with Welch's correction, p<0.0001, n=8 wells per genotype. KOLF: two-tailed unpaired t-test, p<0.0001, n=6 wells per genotype. Graphs represent mean ± SEM. *p≤0.05; **p≤0.01; ***p≤0.001; ****p≤0.0001.

A Uncropped blot from Figure 3B
GIH36C2 donor, Cathepsin D

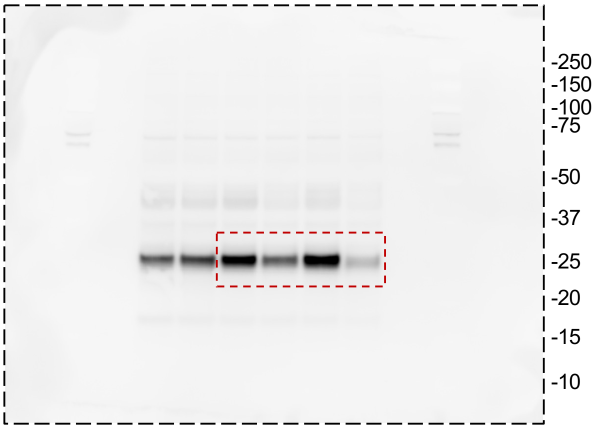


B Uncropped blot from Figure 3B
GIH36C2 donor, Cathepsin B

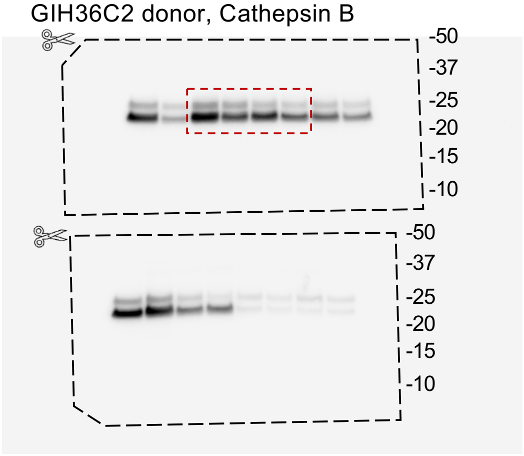


C Uncropped blot from Supp. Figure 4A
GIH178C1 donor, Cathepsin D

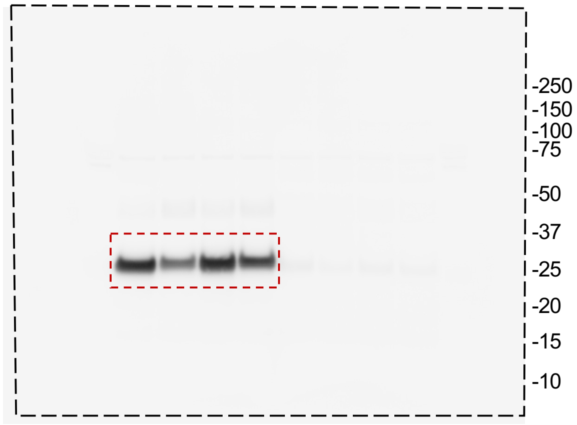


D Uncropped blot from Supp. Figure 4A
KOLF donor, Cathepsin D

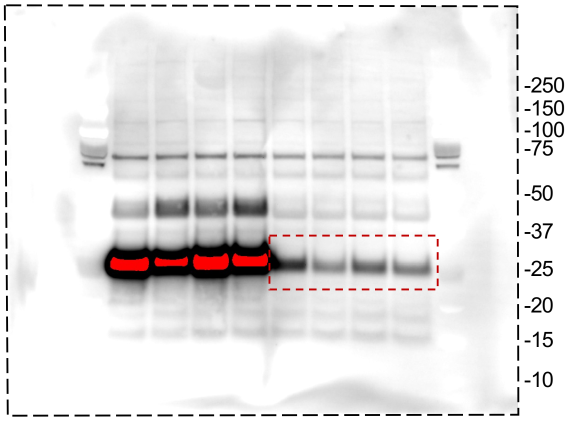


E Uncropped blot from Supp. Figure 4C
GIH178C1 donor, Cathepsin B

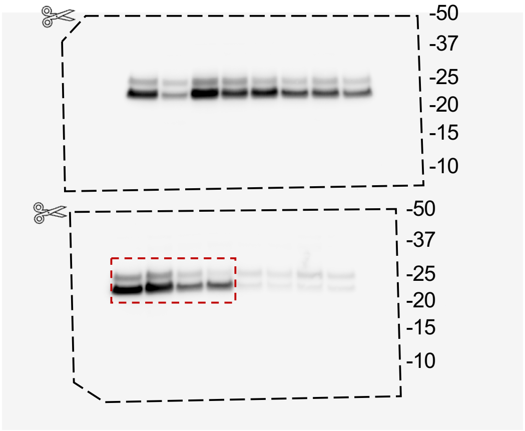


F Uncropped blot from Supp. Figure 4C
KOLF donor, Cathepsin B

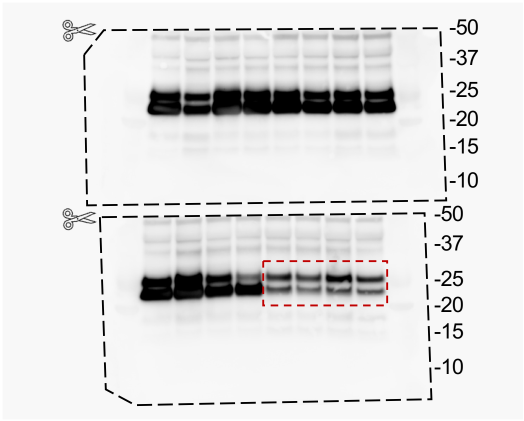

**Supplemental Figure 5. Uncropped immunoblots of cathepsin D and B from in three iPSC donors.** A-F. Uncropped blots. Black dashed line indicates the edge of the membrane. Scissors graphic indicates where membrane was cut prior to antibody incubation. Red dashed line indicates the bands included in representative blots. A. Uncropped blot corresponding to Figure 3B. Lysates from the GIH36 donor were probed for cathepsin D. B. Uncropped blot corresponding to Figure 3B. Lysates from the GIH36 donor were probed for cathepsin B. C. Uncropped blot corresponding to Supplemental Figure 4A. Lysates from the GIH178 donor were probed for cathepsin D. D. Uncropped blot corresponding to Supplemental Figure 4A. Lysates from the KOLF donor were probed for cathepsin D. E. Uncropped blot corresponding to Supplemental Figure 4C. Lysates from GIH178 donor were probed for cathepsin B. F. Uncropped blot corresponding to Supplemental Figure 4C. Lysates from the KOLF donor were probed for cathepsin B. C-D represent the same blot at different exposure times. B and E-F represent the same blot at different exposure times.

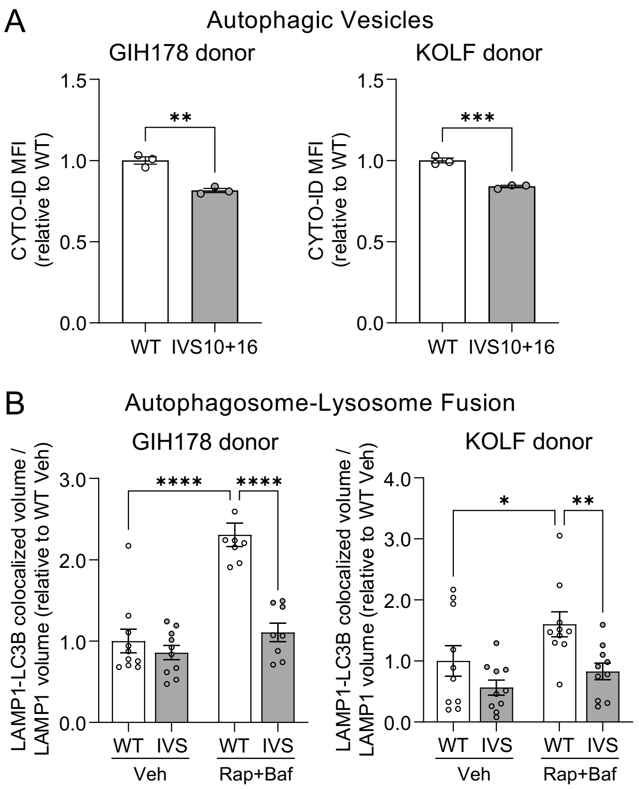


**Supplemental Figure 6. *MAPT* IVS10+16 impairs autophagy by disrupting autophagosome-lysosome fusion in microglia in independent iPSC donors (corresponding to Figure 4).** A. iMG were labeled with CYTO-ID and analyzed by flow cytometry. Quantification of CYTO-ID geometric mean fluorescent intensity (MFI) within the live cell population, expressed relative to WT. n=3 wells per group from one differentiation per donor. Two-tailed unpaired t-test; GIH178, p=0.0019; KOLF, p=0.0007. B. iMG were treated with rapamycin (1µM, Rap) and bafilomycin A1 (400nM, Baf) or DMSO (Veh) and analyzed by immunocytochemistry. Quantification of LAMP1-LC3B co-localized volume was normalized to total LAMP1 volume per field and expressed relative to WT Veh. Ten fields were acquired per group from one differentiation per donor. GIH178: WT Veh, n=30 cells; IVS10+16 Veh, n=31; WT Rap+Baf, n=25; IVS10+16 Rap+Baf, n=25. KOLF: WT Veh, n=17 cells; IVS10+16 Veh, n=20; WT Rap+Baf, n=16; IVS10+16 Rap+Baf, n=17. GIH178 two-way ANOVA: genotype $F(1, 32)=28.68$, $p<0.0001$; treatment $F(1, 32)=38.58$, $p<0.0001$; interaction $F(1, 32)=17.86$, $p=0.0002$. KOLF two-way ANOVA: genotype $F(1, 36)=10.53$, $p=0.0025$; treatment $F(1, 36)=5.394$, $p=0.0260$; interaction $F(1, 36)=0.8227$, $p=0.3704$. P-values by Fisher's LSD test. GIH178: WT Veh vs IVS Veh, p=0.4031; WT Veh vs WT Rap+Baf, p<0.0001; IVS Veh vs IVS Rap+Baf, p=0.1700; WT Rap+Baf vs IVS Rap+Baf, p<0.0001. KOLF: WT Veh vs IVS Veh, p=0.1069; WT Veh vs WT Rap+Baf, p=0.0284; IVS Veh vs IVS Rap+Baf, p=0.3236; WT Rap+Baf vs IVS Rap+Baf, p=0.0058. Graphs represent mean ± SEM. *p≤0.05; **p≤0.01; ***p≤0.001, ****p≤0.0001.

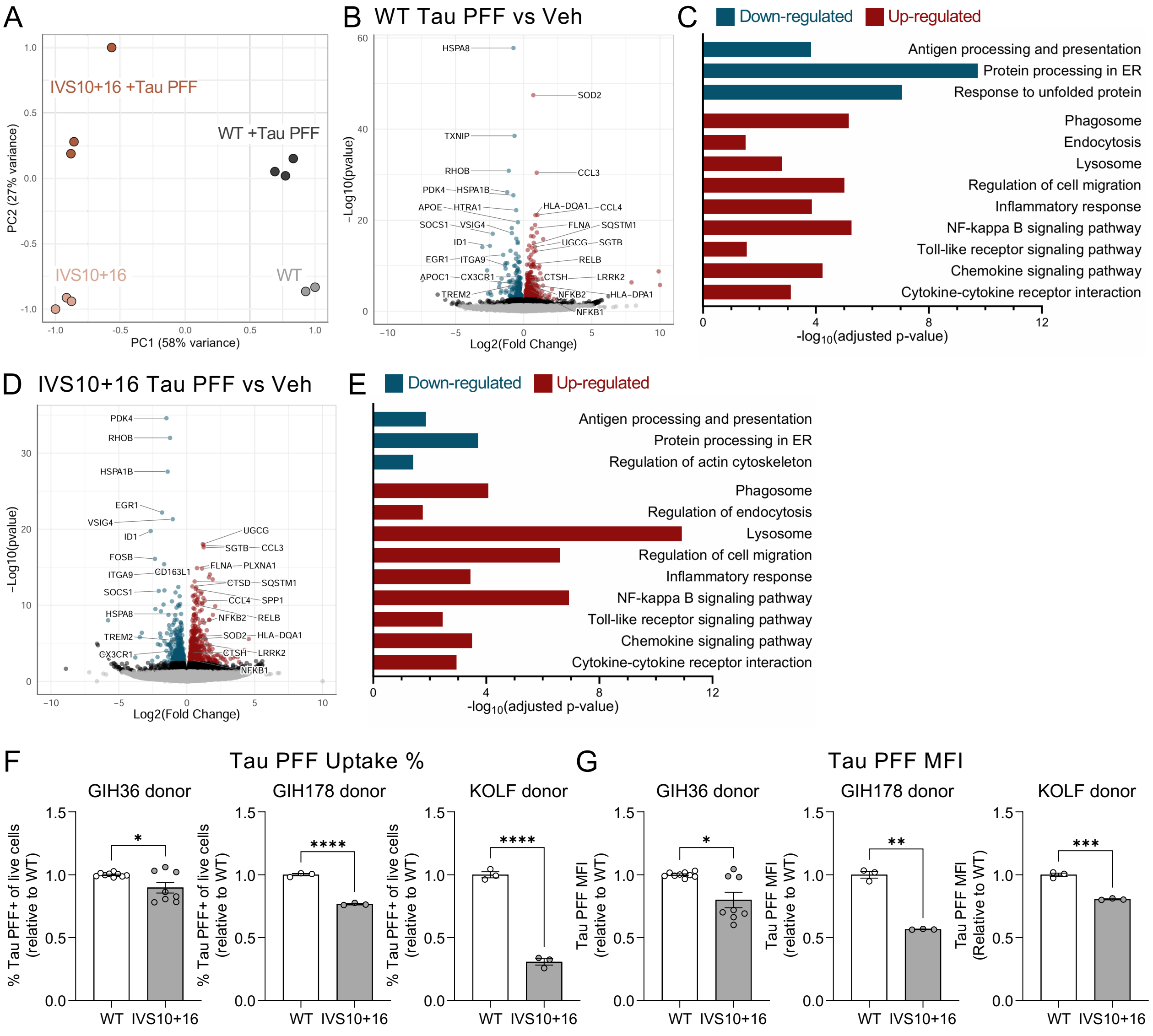

A
IVS10+16 +Tau PFF
WT +Tau PFF
IVS10+16
WT
PC1 (58% variance)
PC2 (27% variance)
B WT Tau PFF vs Veh
HSPA8
SOD2
TXNIP
RHOB
CCL3
PDK4
HSPA1B
APOE
HTRA1
HLA-DQA1
CCL4
SOCS1
VSIG4
FLNA
SQSTM1
ID1
UGCG
SGTB
EGR1
ITGA9
RELB
APOC1
CX3CR1
CTSH
LRRK2
TREM2
NFKB2
HLA-DPA1
NFKB1
Log2(Fold Change)
-Log10(pvalue)
C
Down-regulated
Up-regulated
Antigen processing and presentation
Protein processing in ER
Response to unfolded protein
Phagosome
Endocytosis
Lysosome
Regulation of cell migration
Inflammatory response
NF-kappa B signaling pathway
Toll-like receptor signaling pathway
Chemokine signaling pathway
Cytokine-cytokine receptor interaction
-log10(adjusted p-value)
D IVS10+16 Tau PFF vs Veh
PDK4
RHOB
HSPA1B
EGR1
VSIG4
ID1
UGCG
SGTB
CCL3
FOSB
FLNA
PLXNA1
ITGA9
CD163L1
CTSD
SQSTM1
SOCS1
CCL4
SPP1
HSPA8
NFKB2
RELB
TREM2
SOD2
HLA-DQA1
CX3CR1
CTSH
LRRK2
NFKB1
Log2(Fold Change)
-Log10(pvalue)
E
Down-regulated
Up-regulated
Antigen processing and presentation
Protein processing in ER
Regulation of actin cytoskeleton
Phagosome
Regulation of endocytosis
Lysosome
Regulation of cell migration
Inflammatory response
NF-kappa B signaling pathway
Toll-like receptor signaling pathway
Chemokine signaling pathway
Cytokine-cytokine receptor interaction
-log10(adjusted p-value)
F
Tau PFF Uptake %
GIH36 donor
GIH178 donor
KOLF donor
% Tau PFF+ of live cells (relative to WT)
WT
IVS10+16
G
Tau PFF MFI
GIH36 donor
GIH178 donor
KOLF donor
Tau PFF MFI (relative to WT)
Tau PFF MFI (Relative to WT)
WT
IVS10+16

**Supplemental Figure 7. *MAPT* WT and IVS10+16 iMG upregulate phagosome, lysosome, and inflammatory response pathways after exposure to tau fibrils.** A-E. *MAPT* IVS10+16 and isogenic control iMG were treated with tau PFF (50nM) or equivalent volume DPBS (vehicle) for 24hr and analyzed by RNASeq. Veh-treated samples are the same as in **Figure 1C-E**. WT Veh, n=2; IVS10+16 Veh, n=3; WT Tau PFF, n=3; IVS10+16 Tau PFF, n=3. A. Principal component analyses (PCA) were calculated using the top 500 most variable genes across all groups using rlog-normalized counts in DESeq2. WT Veh, light grey; WT Tau PFF, dark grey; IVS10+16 Veh, light orange; IVS10+16 Tau PFF, dark orange. B-C. Differential gene expression and pathway analysis comparing WT Tau PFF to WT Veh. D-E. Differential gene expression and pathway analysis comparing IVS10+16 Tau PFF to IVS10+16 Veh. B, D. Volcano plots. Blue, significantly down-regulated genes (FDR $p \leq 0.05$; Log2FC<0). Red, significantly up-regulated genes (FDR $p \leq 0.05$; Log2FC>0). Black, $p \leq 0.05$. Grey, not significant. C, E. Pathway analysis of differentially expressed genes (FDR $p \leq 0.05$). Blue bars, down-regulated genes. Red bars, up-regulated genes. F-G. iMG were treated with tau PFF-ATTO488 (500nM) for 24hr and analyzed by flow cytometry. GIH36: Data from three independent differentiations were normalized to WT within each differentiation. WT, n=9 wells, IVS10+16, n=8 wells. GIH178 and KOLF: Data from one differentiation were normalized to WT for each donor. n=3 wells per genotype. F. Quantification of the percentage of live cells which were tau PFF-ATTO488+, normalized to WT. GIH36: two-tailed unpaired t-test with Welch's correction, p=0.0455. GIH178 and KOLF: two-tailed unpaired t-tests, p<0.0001. G. Quantification of ATTO488 geometric mean fluorescent intensity (MFI) within the live cell population, expressed relative to WT. GIH36: two-tailed unpaired t-test with Welch's correction, p=0.0140. GIH178: two-tailed unpaired t-test with Welch's correction, p=0.0037. KOLF: two-tailed unpaired t-test, p=0.0002. Graphs represent mean ± SEM. *$p \leq 0.05$; **$p \leq 0.01$; ***$p \leq 0.001$; ****$p \leq 0.0001$.

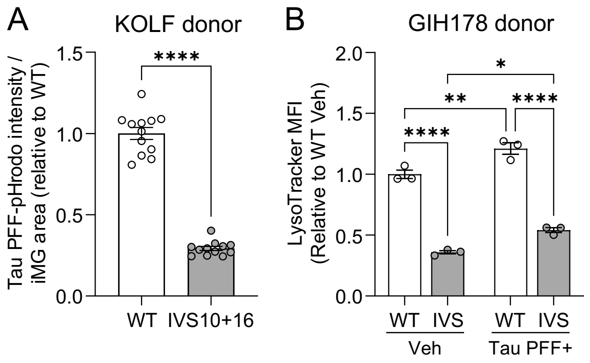


**Supplemental Figure 8. Tau challenge reveals reduced lysosomal capacity in *MAPT* IVS10+16 microglia in independent iPSC donors (corresponding to Figure 5).** A. KOLF donor iMG were treated with tau PFF-pHrodo (250nM) and analyzed by Incucyte live cell imaging. Quantification of tau PFF-pHrodo integrated intensity at 24hrs was normalized to cell area. Data from two independent differentiations were normalized to WT within each differentiation. n=12 wells per genotype. Two-tailed unpaired t-test with Welch's correction, $p<0.0001$. B. GIH178 donor iMG were treated with tau PFF-ATTO488 (500nM) for 24hrs, labeled with LysoTracker for 30mins, and analyzed by flow cytometry. Quantification of LysoTracker geometric mean fluorescent intensity (MFI) within the live cell population for Veh-treated samples and within the Tau PFF+ population for tau-treated samples were normalized to WT Veh. Data from one differentiation with n=3 wells per group. Two-way ANOVA: genotype $F(1, 8)=418.9$, $p<0.0001$; treatment $F(1, 8)=37.70$, $p=0.0003$; interaction $F(1, 8)=0.1978$, $p=0.6683$. Tukey's multiple comparisons test: WT Veh vs IVS Veh, $p<0.0001$; WT Veh vs WT Tau PFF, $p=0.0071$; IVS Veh vs IVS Tau PFF, $p=0.0161$; WT Tau PFF vs IVS Tau PFF, $p<0.0001$. Graphs represent mean ± SEM. *$p\leq0.05$; **$p\leq0.01$; ****$p\leq0.0001$.
